# Graded FGF activity patterns distinct cell types within the apical sensory organ of the sea anemone *Nematostella vectensis*

**DOI:** 10.1101/2023.06.07.544043

**Authors:** Keith Z. Sabin, Shiyuan Chen, Eric Hill, Kyle J. Weaver, Jacob Yonker, MaryEllen Kirkman, Bret Redwine, Anna M. L. Klompen, Xia Zhao, Fengli Guo, Cathy McKinney, Jessica L. Dewey, Matthew C. Gibson

## Abstract

Bilaterian animals have evolved complex sensory organs comprised of distinct cell types that function coordinately to sense the environment. Each sensory unit has a defined architecture built from component cell types, including sensory cells, non-sensory support cells, and dedicated sensory neurons. Whether this characteristic cellular composition is present in the sensory organs of non-bilaterian animals is unknown. Here, we interrogate the cell type composition and gene regulatory networks controlling development of the larval apical sensory organ in the sea anemone *Nematostella vectensis*. Using single cell RNA sequencing and imaging approaches, we reveal two unique cell types in the *Nematostella* apical sensory organ, GABAergic sensory cells and a putative non-sensory support cell population. Further, we identify the paired-like (PRD) homeodomain gene *prd146* as a specific sensory cell marker and show that Prd146^+^ sensory cells become post-mitotic after gastrulation. Genetic loss of function approaches show that Prd146 is essential for apical sensory organ development. Using a candidate gene knockdown approach, we place *prd146* downstream of FGF signaling in the apical sensory organ gene regulatory network. Further, we demonstrate that an aboral FGF activity gradient coordinately regulates the specification of both sensory and support cells. Collectively, these experiments define the genetic basis for apical sensory organ development in a non-bilaterian animal and reveal an unanticipated degree of complexity in a prototypic sensory structure.

## Introduction

The ability of organisms to sense and respond to their environment is essential for their survival. Investigations in select model organisms have provided valuable insights into the development of a few representative sensory systems, such as visual, auditory, and olfactory structures^1–6^. In these cases, sensory structures are generally comprised of three broad cell classes: 1) Ciliated sensory cells that perceive specific environmental inputs (e.g. temperature, light, vibration); 2) Non-sensory cells that support sensory cell function; and 3) Transducing cells, often neurons or the sensory cells themselves, that relay sensory information to modulate organismal behavior. However, known examples only account for a small proportion of the diverse sensory structures present throughout animals and our understanding of the organization, cell type diversity, and developmental mechanisms underlying sensory organ evolution remain incomplete.

A broad diversity of marine invertebrate animals undergo indirect development through planktonic larval forms which are sensitive to different environmental inputs such as light, chemical cues, and mechanical stimuli^7–9^. The perception and integration of these different sensory modalities controls many aspects of larval biology, including swimming, settlement, and metamorphosis^10–14^ . A common anatomical feature found in the larvae of many lophotrochozoans, echinoderms, and hemichordates is a ciliated structure, called the apical sensory organ (**Figure 1A**^15–24)^. This is a prominent larval-specific structure that forms at the apical/aboral/anterior pole of the larval body plan and is adorned with long apical tuft cilia^15, 25^. The apical tuft cilia are generated by apical organ sensory cells and represent a defining feature of apical sensory organs ^25–27^. Apical organ sensory cells have either a columnar or flask-like morphology and generally produce a single apical tuft cilium^16, 18, 21, 25, 26, 28^. In addition, apical organ sensory cells have basally localized nuclei and are surrounded by ectodermal cells with apically localized nuclei. Apical organ sensory cells in the larvae of the annelid *Platynereis dumerilii* and the sea anemone *Nematostella vectensis* express various chemo- and mechano-receptors and express light sensitive proteins, like opsins ^29^ and the *trpV* channel^30^, respectively. Therefore, these cells represent a highly specialized cell type that could be responsive to multiple sensory inputs. In addition to sensory cells, a population of putative larval-specific neurons was found to fully encircle the apical organ sensory cells in *Nematostella* planula larvae^31^. However, the precise molecular identity and developmental mechanisms regulating their development remain unclear. Overall, the exact function of the apical sensory organ is not known but it may be involved in the regulation of settlement and metamorphosis in some animals^10, 13, 23, 32^. Deeper investigations into the function of the apical sensory organ across species will be critical to identify conserved and species-specific aspects of apical organ sensory biology.

**Figure 1:**
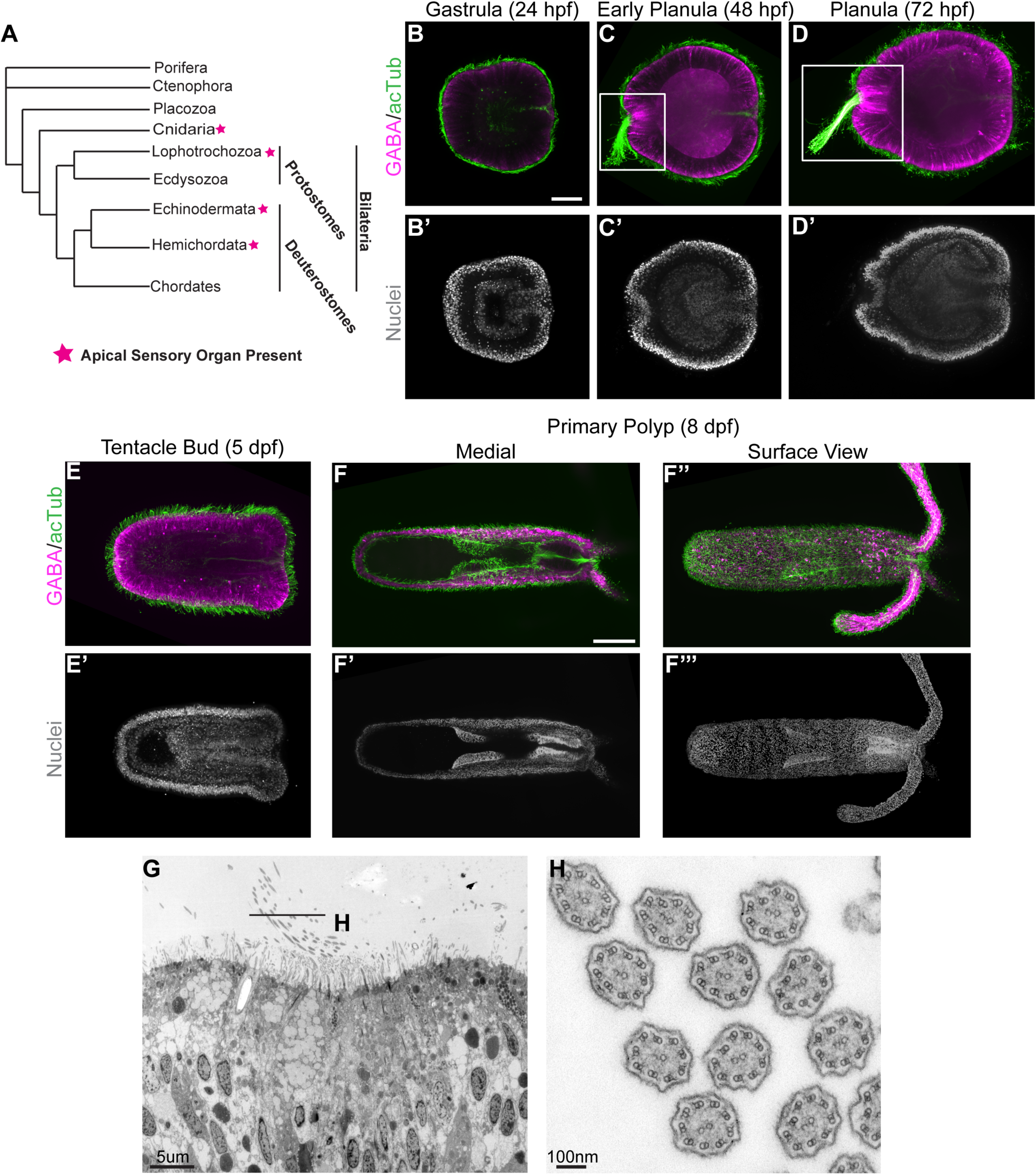
*Nematostella* apical sensory organ development. (A) Schematic phylogeny where animal groups with larvae that have a ciliated apical sensory organ are indicated by a magenta star. (B-F’’’) Fluorescent images showing the general animal morphology (nuclei) and formation of the apical sensory organ (cilia: green/GABA: magenta) at different developmental stages. Scale bar, 50um. (G, H) transmission electron micrographs showing the larval aboral domain (G) and internal axonemal structure of the apical tuft cilia (H). Scale bars 100um (F-F’’’’), 50um (A-E’), 5um (G) and 100nm (H).

In addition to Bilaterian animals (deuterostomes and protostomes), Anthozoan Cnidarians (sea anemones and corals) are the earliest branching metazoans to feature ciliated apical sensory organs, which are found in the planula larvae of actinarian sea anemones (**Figure 1A**^25, 26^). This suggests that anthozoan apical sensory organs represent an ancient structure that was present in the eumetazoan common ancestor over 600 million years ago. Molecular characterization of the larval aboral/apical domain in *Nematostella* (cnidarian), *Platynereis* (protostome), and the sea urchin *Strongylocentrotus purpuratus* (deuterostome) suggest deep molecular conservation^29, 30, 33^. Importantly, the bilaterian anterior markers, *six3* and *foxQ2*, are expressed in the aboral/apical domain of the larvae from these species and the apical organ develops in a region where *six3* and *foxQ2* are repressed or lowly expressed^29, 30, 34^. Further, fibroblast growth factor (FGF) signaling may be broadly required for apical organ development in multiple species since homologues of FGF receptors and ligands are co-expressed in the apical domain of some bilaterian and cnidarian larvae^29, 35, 36^. In *Nematostella*, FGF signaling is responsible for repressing the expression of both *six3/6* and *foxQ2a* in the aboral domain and is required for development of the apical sensory organ^36^. Still, the precise function of FGF signaling during apical sensory organ development is not completely understood.

In *Nematostella* embryos, the expression of an FGF receptor (*fgfra*) and two FGF ligands (*fgfa1, fgfa2*) is initiated by the aboral patterning genes *six3/6* and *foxQ2a* around 11 hours post fertilization (hpf)^35–37^. As development proceeds, expression of the FGF signaling components becomes restricted to the apical sensory organ and forms a positive feedback loop which, in part, prevents its degeneration^35, 36^. Binding of the FGF ligands to FGFRa results in the activation of a MAP kinase cascade and phosphorylation of the extracellular regulated kinase (ERK)^35^. Activation of this pathway induces the aboral expression of key effector transcription factors, Hox1 and SoxB1, which are critical for aboral patterning and apical organ development^35, 36^. FGFR-MAPK signaling specifies both apical organ sensory cells and the putative larval-specific neurons that encircle the apical organ^31, 35, 36^. The precise mechanisms by which FGF signaling specifies two distinct cell types during apical organ development is not clear. During vertebrate development, FGF signaling can function in a classical morphogen gradient fashion to specify unique cell identities in a concentration dependent manner^38–40^. Whether this form of a classic FGF gradient is utilized during the development of earlier branching animals is not known.

Here, we took an unbiased single cell RNA sequencing-based approach (scRNA-seq) to characterize the apical sensory organs of *Nematostella* larvae. Our analysis identified two major cell types: sensory cells and a ring of cells that encircles them. Subsequent functional experiments revealed that the cells encircling the apical organ are not likely to be neurons. Intriguingly, the anatomical organization of these cell types is similar to that observed in the teleost neuromast, with centrally located sensory cells surrounded by non-sensory support cells (mantle cells)^41, 42^. Therefore, we proposed that the cells encircling the apical organ sensory cells are a type of support cell. We further identified a paired-type (PRD) homeodomain-containing transcription factor, Prd146, that is specifically expressed within the apical organ sensory cells. Mutagenesis of *prd146* with CRISPR/Cas9 revealed that Prd146 is required for apical sensory organ development and that *prd146* mutant larvae successfully transitioned into primary polyps. Mechanistically, Prd146 is a key regulator of the apical sensory organ gene regulatory network downstream of FGF signaling. Finally, we show that an FGF signaling gradient is required to specify both sensory cell and support cell fates during apical sensory organ development. High levels of FGF signaling specifically induce *prd146* expression, which in turn functions to specify sensory cell and repress support cell fates. Taken together, these results provide key resources for future studies of the *Nematostella* apical sensory organ and reveal an unanticipated degree of cellular and molecular complexity within a prototypic sensory structure.

## Results

### Characterizing apical sensory organ development in Nematostella larvae

Prior to interrogating cellular diversity within the apical sensory organ, we first generated a time course of *Nematostella* development at 24°C. *Nematostella* development is rapid, progressing from the gastrula stage at 24hpf (**Figure 1B’**) to motile planula larvae by 48-72hpf (**Figure 1C’, D’**). By 5dpf, larvae have largely settled and transitioned into the tentacle bud stage (**Figure 1E’**), followed by the primary polyp stage at 8dpf (**Figure 1F’, F’’’**). Apical organ sensory cells and the apical tuft cilia can be visualized using antibodies against the neurotransmitter GABA and acetylated tubulin, respectively^35, 43^. This analysis revealed no apical tuft cilia and very few GABA^+^ cells present in gastrulae (24hpf); only motile cilia were present on ectodermal cells (**Figure 1B**). However, by early larval stages (48hpf), we saw an accumulation of GABA^+^ cells at the aboral pole, corresponding to apical organ sensory cells (**Figure 1C; white box**). Importantly, those cells were also beginning to generate long apical tuft cilia (**Figure 1C; white box**). The GABAergic sensory cells and apical tuft cilia persisted through the planula stage (**Figure D; white box**) but were largely absent from tentacle bud stage animals and primary polyps (**Figure E, F, F’**). Other GABA^+^ cells, which likely represent neural cell types^43, 44^, were observed along the oral-aboral axis in planula larvae, tentacle bud, and primary polyp stages (**Figure 1C, D, E, F, F’’**). Transmission electron micrographs through the aboral domain clearly showed the long apical tuft cilia that emanated from apical organ sensory cells (**Figure 1G**). Cross sections through the apical tuft cilia revealed a clear 9+2 axonemal structure (**Figure 1H**).

### Mapping cell type diversity in the larval aboral pole and apical sensory organ

To investigate the cell type composition of apical sensory organs, we generated a 10x single cell RNA sequencing data set of whole *Nematostella* planula larvae at 72hpf, a time point where we know the apical organ is fully formed (**Figure 1D**). Subsequent clustering analysis detected known larval cell types, supporting the overall accuracy of our data set (**Figure S1, Supplemental Table 1, 2**)^45, 46^. Importantly, apical organ sensory cells, which express *fgfa1*, formed their own unique cluster (**Figure S1**). To further characterize cell types associated with the aboral domain and the apical sensory organ, we next subclustered cells that expressed the aboral marker genes *six3/6* and *foxQ2a*^36^ as well as cells that comprised the apical organ cluster (**Figure S2A**). This approach revealed seven transcriptionally distinct cell clusters which correspond to six distinct cell types within the aboral domain (**Figure 2A**). Based on the expression of known and novel marker genes, we determined that four of the six cell clusters represented cell types not restricted to the apical organ. These included neurons and neurosecretory cells (*elav, insm1, cd151*), progenitor cells (*soxC*, *soxB2, cdk1*), embryonic ectoderm (*glycoprotein 1*), and cnidocytes (*nematogalectin*, *rhamnose-binding lectin*), (**Figure 2B-E’, Figure S2B, Supplemental Table 3**). Our analysis identified two cell types that were unique to the apical sensory organ: the sensory cells themselves (*fgfa2, prd146*) (**Figure 2F, F’, Figure S2B**) and a population of cells that encircle the sensory cells, hereafter referred to as support cells (*poxA, slc26a6, amt1*) (**Figure 2G, G’; green box, Figure S2B**). Many of the known apical organ/aboral marker genes in *Nematostella* were expressed in both the sensory cells and support cells (*six3/6*, *hox1*, *soxB1*) (**Figure S3A**,**B,D,E**). However, *foxQ2a* was uniquely expressed in support cells but not the sensory cells (**Fig. S3C**), while a paired-like (PRD) homeodomain containing transcription factor was the only transcription factor analyzed that was expressed only in the sensory cells (**Figure 2F, F’, Figure S3F**). This PRD-like gene was the most highly expressed marker gene in the sensory cell cluster (**Figure S2B**, **Supplemental Table 3**) and had been previously annotated as *aristaless-like 1* (*alx1*)^47^. Deeper bioinformatic analysis revealed that this locus has been variously identified as *NvHD146*^48^, *Q50-6*^49^, *prd146*^50^, and *ISX-like*^31^. Collectively, these phylogenetic studies and our own maximum likelihood reconstruction of select PRD genes (**Figure S4, Supplemental Table 4**) firmly placed this gene within the PRD-like family of homeodomain genes. However, given the relatively poor statistical support of this gene within a specific gene family (**Figure S4**)^31, 48–51^, we will refer to this gene as *prd146*^50^. Interestingly, and consistent with a recent report^31^, we found that *prd146* formed a well-supported (bootstrap = 98) orthologous group with three additional anthozoan sequences suggesting that this gene may be anthozoan-specific (**Figure S4**).

**Figure 2:**
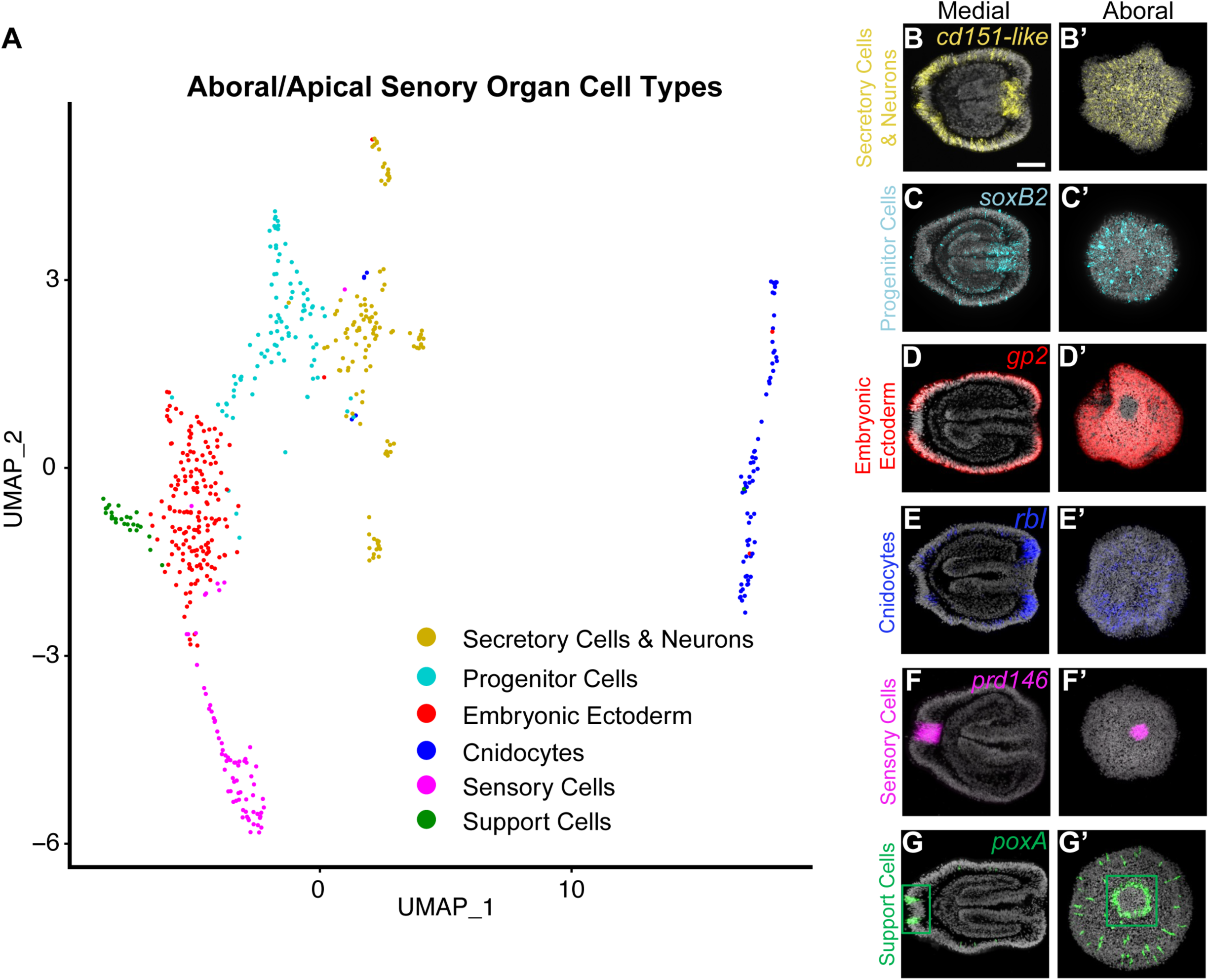
Identification of apical sensory organ-specific cell types. (A) UMAP representation of unique cell types present in the aboral domain and apical sensory organ of *Nematostella* larvae. (C-G’) Representative images of fluorescent *in situ* hybridizations against specific marker genes used to visualize aboral/apical sensory organ cell types. At least 10 larvae were imaged per marker gene. FISH images were pseudo colored to correspond with the cell type indicated in the UMAP (A). Scale bar is 50um.

### Prd146^+^ cells are post-mitotic in larval stages

Our single cell RNA sequencing analysis identified *prd146* as a specific apical organ sensory cell marker (**Figure 2F, F’, Figure S3F**). To better study the development of this cell population, we generated polyclonal anti-Prd146 antibodies and performed a time course analysis. In gastrulae (24hpf), Prd146 was broadly expressed within the aboral ectoderm (**Figure S5A**) but became restricted to the apical sensory organ during early to late planula stages (48-96hpf) (**Figure S5B-D**). Prd146 expression was largely absent from tentacle bud stage animals and primary polyps (**Figure S5E-F’**), which is consistent with the apical sensory organ being a larval-specific structure (**Figure 1B, C, D, E, F**). Further, these results corroborate the expression dynamics of the *prd146* mRNA^31^.

Within diverse animal sensory organs, mature sensory cells are often post-mitotic. We therefore tested whether Prd146^+^ cells are post-mitotic in *Nematostella* embryos and larvae. In gastrula stage embryos pulsed with EdU, approximately 55% of Prd146^+^ cells were also EdU^+^, showing that these cells are proliferative (**Figure 3 A-C’’, P**). However, when either 48hpf, 72hpf, or 96hpf planulae were pulsed with EdU, there were very few Prd146^+^/EdU^+^ cells showing that they had become post-mitotic (**Figure 3 D-L’’, P**). To determine if this merely reflected a general decrease in proliferation, we quantified the proportion of all larval cells that were also EdU^+^. While there was a gradual decrease in the proportion of EdU^+^ cells as development progressed, the decrease in cell proliferation was not as dramatic as observed in Prd146^+^ cells (**Figure 3Q**). Furthermore, once Prd146^+^ cells became post-mitotic, the average number of Prd146^+^ cells did not increase (**Figure 3R**), indicating that most apical organ sensory cells differentiate by early larval stages (48hpf). Collectively, these results show that Prd146-expressing cells proliferate during early development. As development proceeds, these cells become post-mitotic and exist the cell cycle between 24-48hpf.

**Figure 3:**
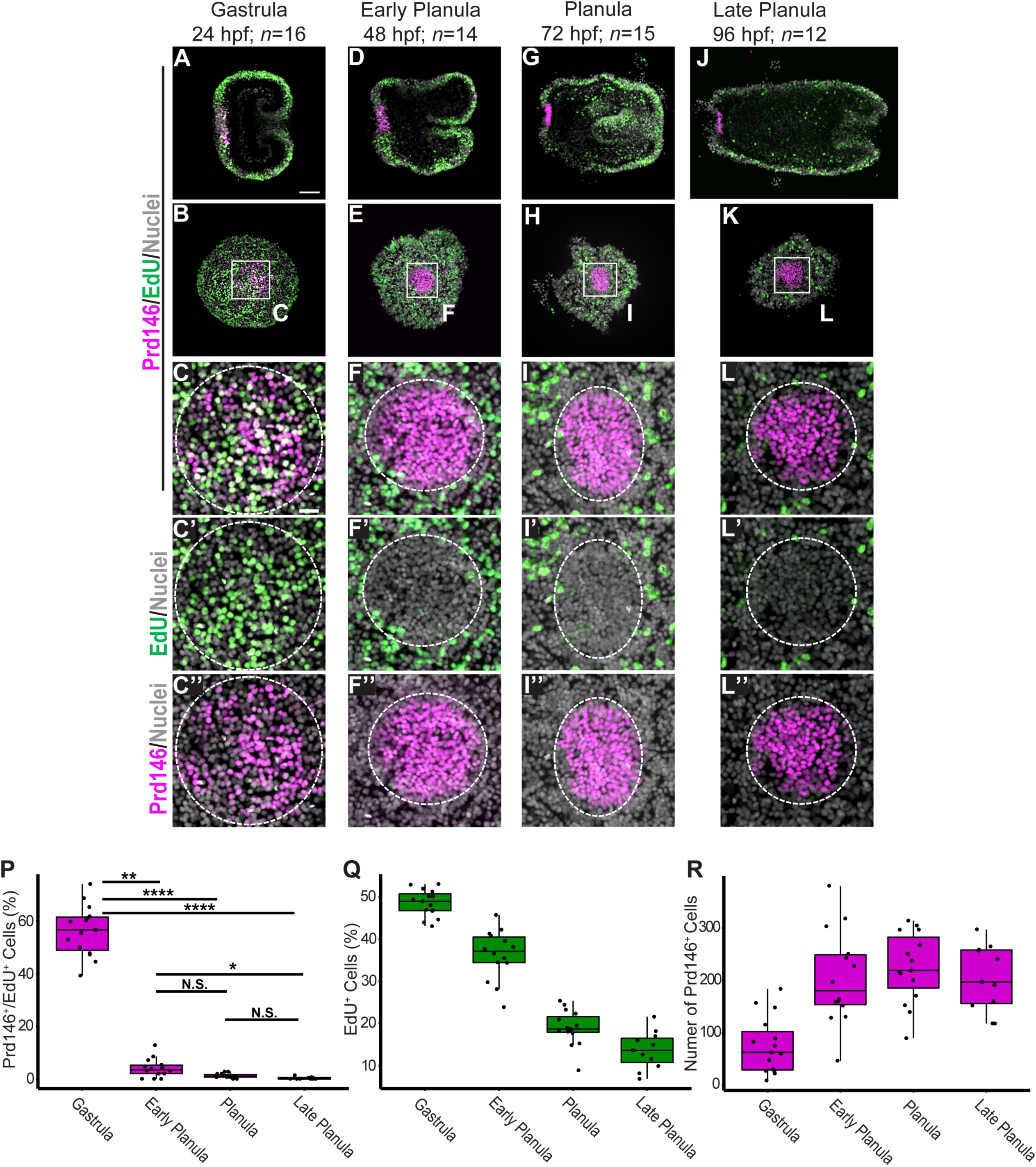
Prd146^+^ cells are post mitotic at larval stages. (A-L’’) Gastrula and larvae were pulsed with EdU (green) for 30 minutes at the indicated developmental stage before being fixed and then co-stained for Prd146 (magenta). (P) Quantification of the average number of Prd146^+^ cells at gastrula and larval stages. (Q) Quantification of the percent of all cells that were EdU^+^ at the indicated time points. (R) Quantification of the proportion of Prd146^+^ cells that were also EdU^+^. Initial differences among groups was determined by the non-parametric Kruskal-Wallis test (*H*(2) = 40.6, *p <* 0.0001, 17^2^_H_ = 0.74). Significant differences between the groups was determined using Dunn’s post hoc test. *p.adjust* < 0.0001 ****, *p.adjust* < 0.01 **, *p.adjust* < 0.05 *, N.S. is “not significant”. Scale bar, 50um.

### Prd146 drives apical organ development downstream of FGF signaling

The spatially restricted expression of *prd146* suggests that it could play a crucial role in regulating apical sensory organ development. Indeed, *prd146* was recently shown to be required for the formation of the apical tuft cilia^31^. To determine whether *prd146* is more broadly required for the specification of apical organ sensory cells, we performed knockdown experiments with two independent shRNAs followed by GABA immunostaining to visualize apical organ sensory cells. Knockdown of *prd146* resulted in a complete loss of Prd146 protein as well as GABAergic sensory cells and the associated apical tuft cilia (**Figure S5G-L**). We next utilized CRISPR/Cas9-mediated mutagenesis to establish a stable F1 *prd146* mutant line that was heterozygous for a +4bp (*prd146^+^*^4^) insertion. The insertion resulted in a frameshift that caused a premature stop codon, truncating the protein prior to the homeodomain. Subsequent analysis confirmed that *wild-type* and heterozygous planulae displayed normal Prd146 expression while homozygous mutants completely lacked Prd146 protein (**Figure 4A-C**). Similarly, both wild type and heterozygous planulae developed normal apical sensory organs while homozygous mutant planulae lacked both GABAergic sensory cells and the apical tuft cilia (**Figure 4D-F**). These findings show that Prd146 is essential for development of the apical sensory organ.

**Figure 4:**
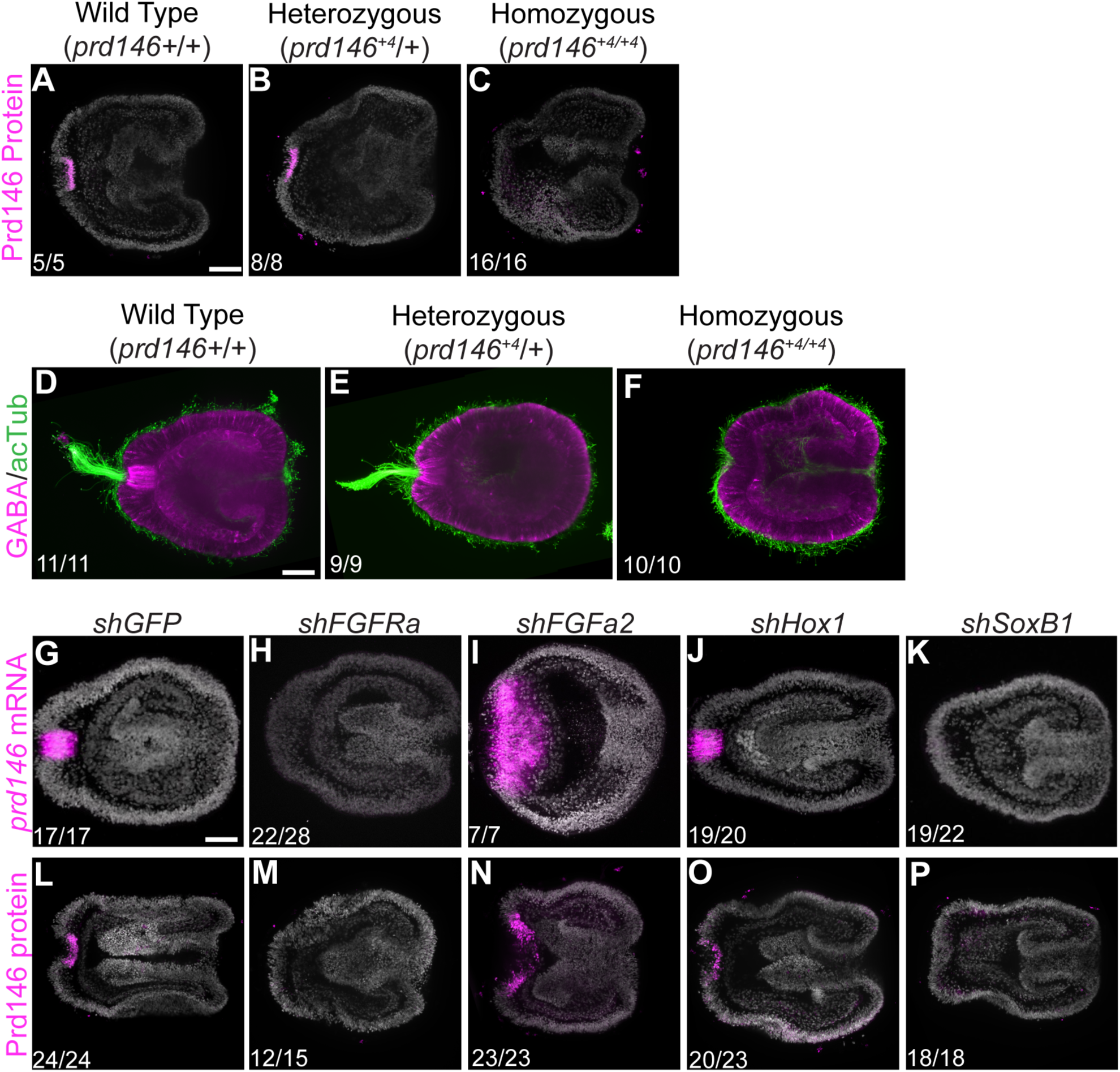
Prd146 is required for apical sensory organ development and is a SoxB1 target gene downstream of FGF signaling. (A-C) Immunofluorescent images of Prd146 protein expression (magenta) in wild type (*prd146* +/+), heterozygous (*prd146^+^*^4^/+), and homozygous mutant (*prd146^+^*^4^/prd146*^+^*^4^) sibling planulae. (D-F) Representative images showing apical sensory organ development in wild type (*prd146* +/+), heterozygous (*prd146^+^*^4^/+), and homozygous mutant (*prd146^+^*^4^/prd146*^+^*^4^) sibling planulae. Expression of *prd146* transcript (G-K) and protein (L-P) is normal in control larvae (G, L), after knockdown of *fgfra* (H, M), *fgfa2* (I, N), *hox1* (J, K), and *soxB1* (K, P). Scale bar, 50um.

To test whether the apical sensory organ is required for the larval to polyp transition in *Nematostella*, we reared progeny of an F^1^ *prd146^+^*^4^/+ in-cross until 9dpf and observed that essentially all planulae had successfully transitioned into primary polyps. We genotyped 92 randomly selected polyps from 2 independent spawning events and found a Mendelian ratio of the expected genotypes (wild type: 21/92 (22.8%, heterozygous: 49/92 (53.2%), homozygous: 23.9% (22/92)) (**Figure S5M**). We conclude that *prd146* mutant planulae lacking an apical sensory organ can successfully transition into primary polyps under laboratory conditions. Closer inspection of 9dpf primary polyps revealed normal morphology in wild type, heterozygous, and homozygous mutant polyps (**Figure S5N-P**). Collectively, these results show that Prd146 is required for apical organ development but dispensable for the larval-to-polyp transition in *Nematostella* under laboratory conditions. However, this does not rule out the possibility that the apical organ could play a role in controlling the timing or other aspects of the larval-polyp transition under environmental conditions.

Next, we investigated potential upstream signals that could drive the spatially restricted pattern of *prd146* expression during larval development. Consistent with a previous report^37^, knockdown of *fgfra* led to a complete loss of *prd146* transcript (**Figure 4H compared to 4G**). Leveraging our anti-Prd146 antibody, we confirmed that *fgfra* knockdown results in a corresponding loss of Prd146 protein (**Figure 4M compared to Figure 4L**). Further, shRNA-mediated knockdown of FGFa2 ligand, which normally restricts apical organ size^35^, led to a dramatic increase in the Prd146 expression domain (**Figure 4I, N**). In *Nematostella*, Hox1 and SoxB1 are key effector transcription factors downstream of FGF signaling that are required for aboral patterning and apical organ development^36^. We found that shRNA-mediated knockdown of *soxB1* but not *hox1* resulted in a loss of Prd146 expression (**Figure 4J-P**), consistent with the non-overlapping functions of SoxB1 and Hox1^36^. Taken together, these results identify a previously unappreciated role for Prd146 during the specification of apical organ sensory cells. In addition, our data show that *prd146* is a SoxB1 target gene downstream of aboral FGF signaling. Collectively, these findings define key components of the gene regulatory network required for apical organ development in *Nematostella*.

### PRD146 is a central node in the apical organ gene regulatory network

While the experiments above demonstrate that Prd146 is required for sensory cell specification, the downstream mechanisms of action remain unclear. We thus sought to identify Prd146 target genes involved in aboral patterning and apical organ development^35, 36, 52, 53^. We first examined the effect of Prd146 on the expression of the aboral patterning genes *six3/6* and *foxQ2a*, which are normally expressed throughout the aboral ectoderm in gastrula but then specifically repressed in the apical sensory organ of planula larvae^36^. Following knockdown of *prd146*, *six3/6* and *foxQ2a* expression failed to be repressed within the apical sensory organ (**Figure 5A-D**), suggesting that Prd146 represses both *six3/6* and *foxQ2a*. Because *prd146* is an FGFRa target gene (**Figure 4H, M**), we also explored whether Prd146 in turn regulates the expression of *fgfra* to form a regulatory feedback loop. Indeed, knockdown of *prd146* resulted in decreased expression of *fgfra* within the aboral pole while endodermal expression of *fgfra* was unaffected (**Figure 5E, F**). In addition to *fgfra*, knockdown of *prd146* resulted in a loss of *fgfa2*, *foxJ1*, and *coe* expression within the aboral pole (**Figure 5G-L**) while *fgfa1*, *hox1*, and *soxB1* were not affected (**Figure 5M-R**). Together, these results show that Prd146 is a key regulator or the apical organ gene regulatory network (**Figure 5S**).

**Figure 5:**
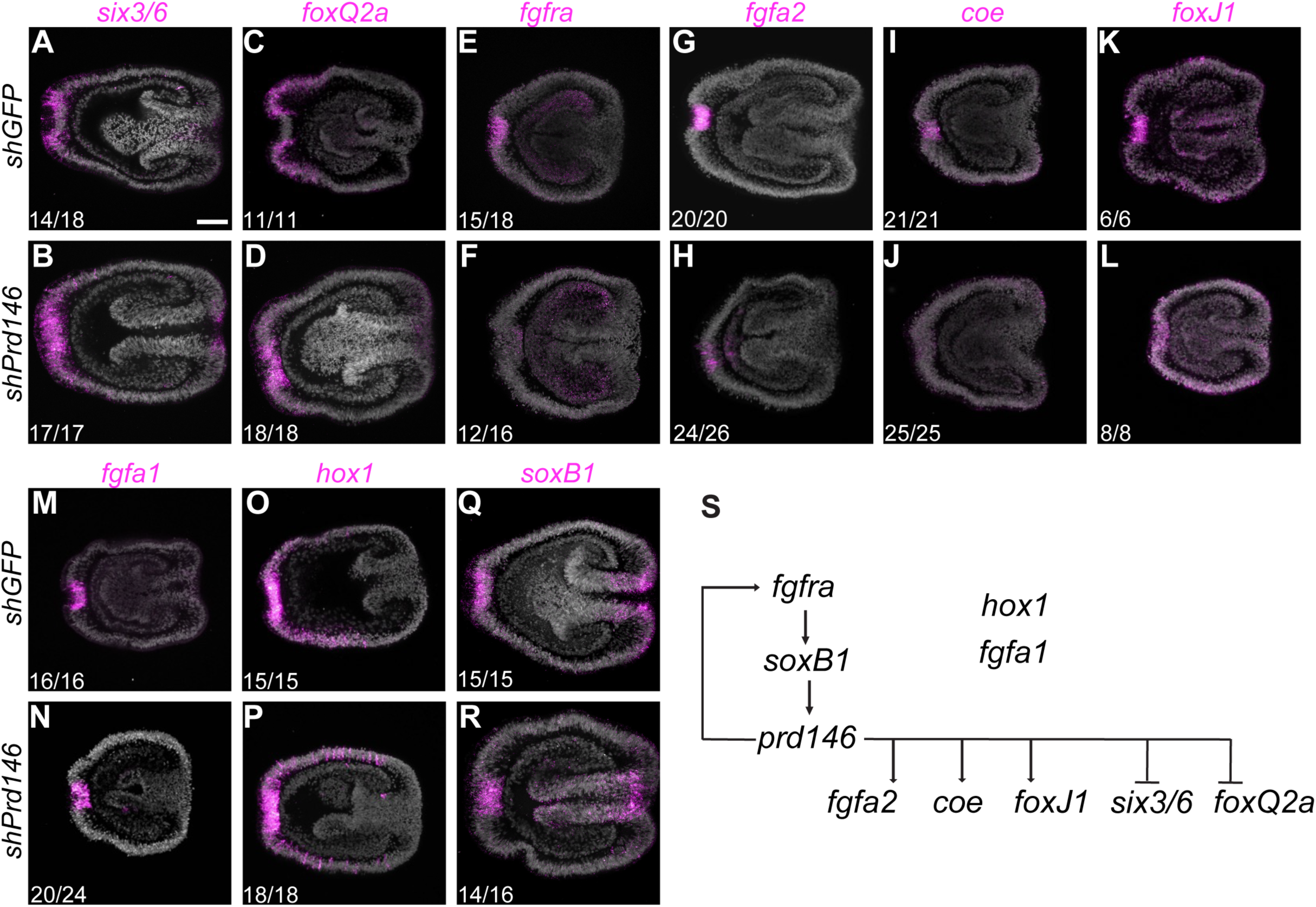
Prd146 is a key regulatory of the apical organ gene regulatory network. (A-R) Fluorescent in situ hybridization against key genes involved with aboral patterning and apical organ development in control and *prd146* knockdown larvae. Prd146 represses expression of the aboral patterning genes *six3/6* (A, B) and *foxQ2a* (C, D), is required for the expression of *fgfra* (E, F), *fgfa2* (G, H), *coe* (I, J), and *foxJ1* (K, L), and *prd146* knockdown does not affect expression of *fgfa1* (M, N), *hox1* (O, P), or *soxB1* (Q, R). (S) Schematic summary of the upstream regulators and downstream target genes of Prd146.

### Molecular characterization of apical sensory organ support cells

In addition to characterizing the sensory cell lineage, we next sought to molecularly define the apical sensory organ support cells. We first performed *in situ* hybridization against two marker genes that our scRNA-seq clustering analysis identified as highly enriched within the support cell cluster: the peroxidase-domain containing gene *peroxinectin A* (*poxA*), and the solute carrier encoded by *slc26a6*. Both genes were expressed by a population of cells that formed a ring around the apical organ sensory cells as well as a cell population found more diffusely throughout the larval body column (**Figure 2G,G’, S6A-B’**). Recently, cells that express *poxA* or *slc26a6* were described as larval-specific neurons based on comparisons between aborally-enriched bulk RNA-seq data and a planula single cell RNA-seq data set^31, 46^. Whether all cells that express these marker genes represent the same cell population and whether these cells are truly neurons has not been experimentally verified. To explore this further, we first examined the expression of *poxA* and *slc26a6* in our data set and found that these genes were largely co-expressed in the same cells (**Figure S6C; black boxes**). Due to their striking overlap and the strength and specificity of the *poxA* in situ probe, we focused our subsequent analysis on *poxA^+^* cells.

While the expression of *poxA* was enriched in the support cell cluster, it was also expressed by a subset of cells in the secretory cells & neurons cluster (**Figure S7A; black boxes**). The *poxA^+^* cells in the secretory cells & neurons cluster co-expressed the neuronal marker *elav*, suggesting that these cells are indeed neurons (**Figure S7B; black box**). Importantly, *elav* was not expressed in the support cell cluster nor was any other known neuronal marker. During *Nematostella* neural development, inhibition of Notch signaling results in an expansion of neuronal cells^54, 55^ and variably affects glad/secretory cell development^56^. It follows that if all *poxA*^+^ cells are larval-specific neurons, then Notch inhibition should elicit an expansion of the cells around the apical organ. To test this, we inhibited Notch signaling by treatment of gastrula to planula stage animals with DAPT. As expected, this resulted in an expansion of the diffuse population of *poxA^+^* cells within the larval body column (**Figure S7C-F; white arrow heads**). In contrast, we observed a complete loss of the population *poxA^+^* cells that ring the apical organ (**Figure S7C-F; white box**). We next repeated these experiments but used a support cell specific marker, *ammonium transporter 1* (*amt1*), identified by our scRNA-seq clustering analysis (**Figure S7G; black box**). Importantly, *amt1* was co-expressed in *poxA*^+^ cells in the support cell cluster but not in secretory cells and neurons (**Figure S7H; black box**). Inhibition of Notch signaling led to a complete loss of *amt1^+^* cells surrounding the apical organ, consistent with our results looking at *poxA^+^* cells (**Figure S7I-L**). Collectively, these results suggest that the lateral population of *poxA* expressing cells are larval-specific neurons and that the *poxA*^+^ cells encircling the apical organ sensory cells are a distinct cell type. We propose that this population represents a non-sensory support cell type similar to that found in other sensory organs in diverse organisms^57–59^.

### A complex transcription factor network controls sensory and support cell specification

Relative to apical organ sensory cells, very little is known about the molecular mechanisms that regulate support cell specification. Bioinformatic analysis indicates that *prd146* is specifically expressed in sensory cells while *hox1* and *soxB1* are expressed in both sensory and support cells (**Figure S3**). We therefore tested the function of these transcription factors during support cell development (**Figure. 6A**). Given that *prd146* is only expressed in sensory cells, we reasoned that *prd146* may repress support cell specification. Confirming this, knockdown of *prd146* resulted in an expansion of *poxA^+^* support cells throughout the aboral pole (**Figure 6C, C’ compared to 6B, B’, K**). Similarly, *soxB1* knockdown resulted in an expansion of support cells throughout the aboral pole (**Figure 6D, D’, K**). In contrast, *hox1* knockdown inhibited or completely blocked the development of support cells, suggesting that Hox1 is required for support cell specification (**Figure 6E-H’, K**). Considering that Hox1 functions in parallel to SoxB1 and Prd146 (**Figure 4J, O, Figure 6A**)^36^, we hypothesized that the expansion of support cells after *soxB1* or *prd146* knockdown likely requires Hox1 function. Indeed, combinatorial knockdown of either *hox1* and *prd146* or *hox1* and *soxB1* inhibited or blocked support cell specification, similar to *hox1* knockdown alone (**Figure 6F-J’, K**). Collectively, these experiments demonstrate that SoxB1 and Prd146 repress support cell specification while Hox1 independently promotes their development (**Figure 6L**).

**Figure 6:**
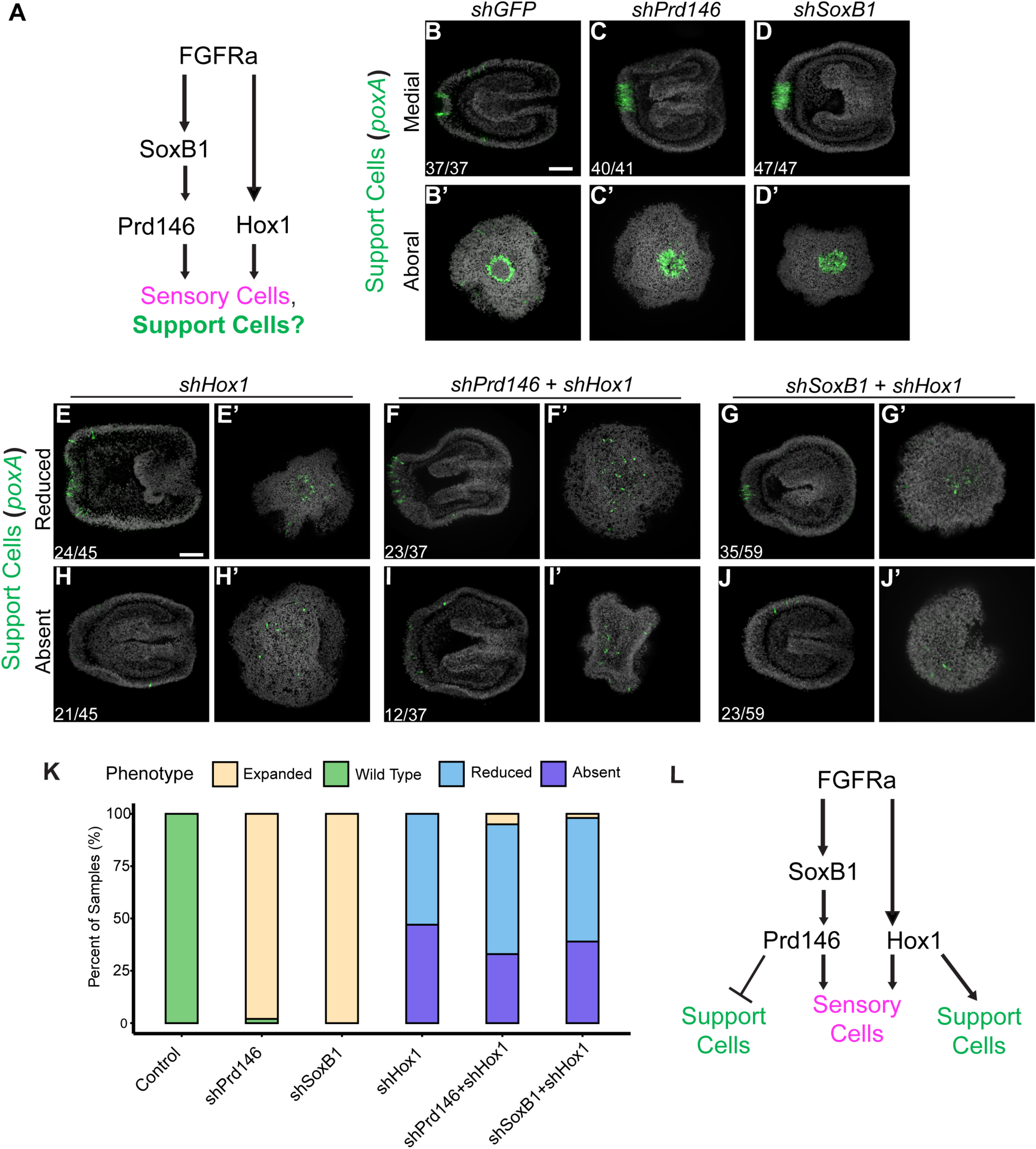
A complex transcription factor network controls support cell specification. (A) Schematic diagram of how FGF signaling regulates sensory cell and maybe support cell specification. (B-D’) Fluorescent in situ hybridization images showing the expression of the support cell marker poxA in control larvae (B, B’) and after knockdown of *prd146* (C, C’), *soxB1* (D, D’), *hox1* (E-H’), *prd146*+*hox1* (F-I’), and *soxB1*+*hox1* (G-J’). (K) Qualitative assessment of the different support cell phenotypes after single or double gene knockdown. (L) Schematic diagram depicting the roles of key FGF target genes regulate sensory and support cell specification. Scale bar, 50um.

### An FGF activity gradient specifies both sensory and support cell lineages

FGF signaling is required for both sensory and support cell specification (**Figure 4H, M**)^31, 35^. Precisely how FGF signaling induces two distinct cell types is not clear. Inhibition of FGF signaling by shRNA-mediated knockdown of *fgfra* resulted in a complete loss of *poxA^+^* support cells from the aboral domain in about 80% of injected animals (**Figure 7A, B**; *n*= 25/31). Intriguingly, *fgfra* knockdown did not affect the development of *poxA* expressing neurons within the larval body column (**Figure 7A-B; white arrowheads**). This further supports our conclusion that these cells represent a distinct population. Remarkably, in 20% of *fgfra* knockdown planulae, we observed aberrant *poxA* expression in the aboral-most cells, which normally develop into sensory cells (**Figure 6C**; *n*= 6/31). Together, these observations suggest that an FGF activity gradient centered on the aboral pole could coordinately regulate both sensory and support cell specification, with high levels of FGF driving sensory identity and low levels driving support identity. To test this, we treated gastrula-to planula-stage animals with increasing concentrations of the FGFR inhibitor SU5402^60^ (**Figure 7D**). This approach inhibits FGF signaling activity in a dose-dependent manner^61^ and allowed us to assay how different levels of FGF activity affected sensory and support cell specification. Treatment of developing animals at the lowest concentration of SU5402 dramatically reduced the size of the *prd146* expression domain with a concomitant decrease in the support cell ring circumference compared to DMSO-treated controls (**Figure 7G, G’ compared to E, E’**). At a slightly higher concentration of FGFR inhibitor, we observed a complete loss of *prd146* expression while cells at the aboral most tip of the planula expressed the support cell marker *poxA* (**Figure 7I, I’**). Finally, at the highest concentration of FGFR inhibitor, which eliminates FGF signaling activity^35, 62^ we observed a complete loss of both sensory and support cells (**Figure 7K, K’**). These results suggest a model where an aboral gradient of FGF signaling coordinately specifies both sensory and support cells fates in a concentration dependent manner (**Figure 7L**).

**Figure 7:**
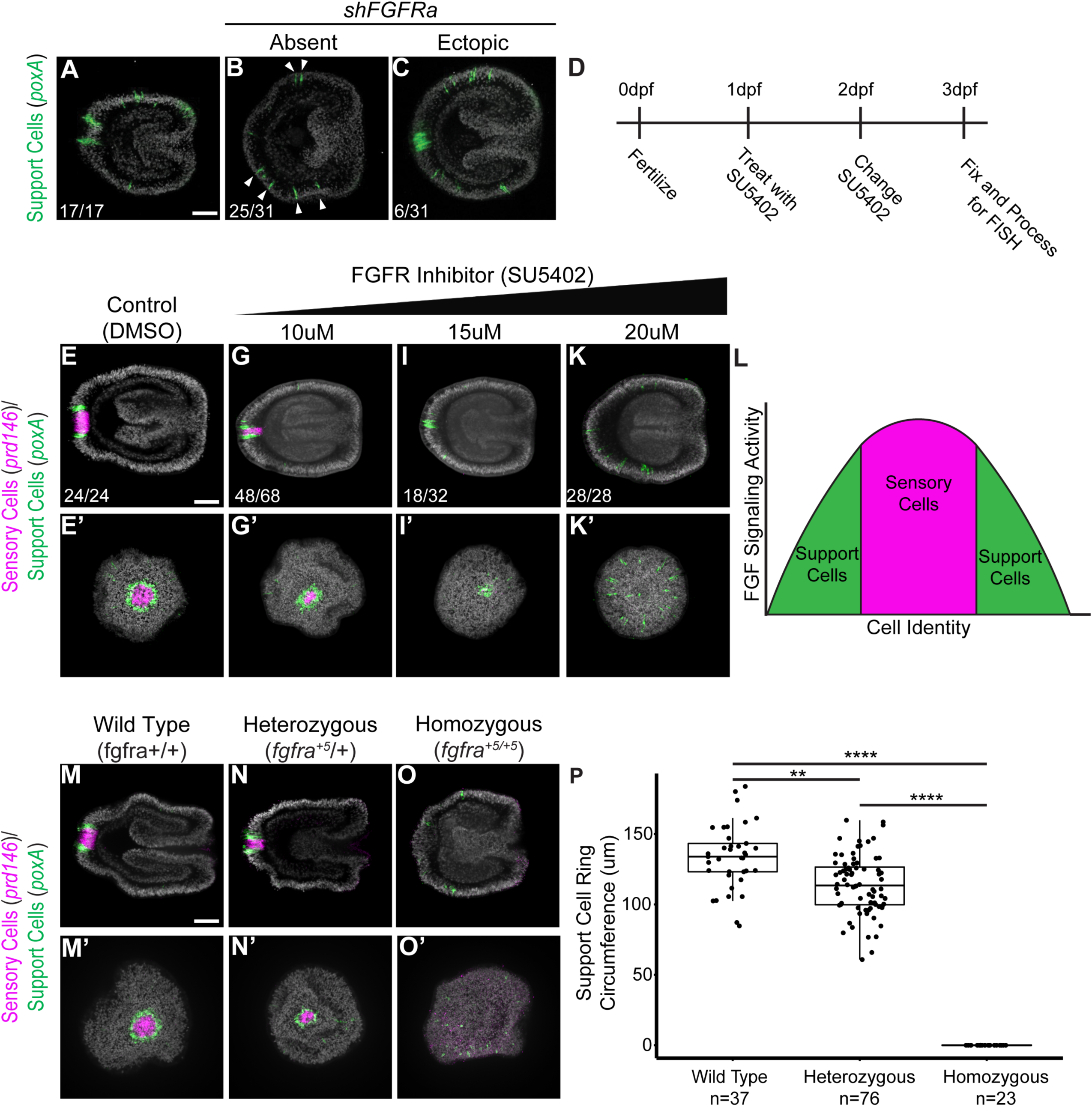
An FGF activity gradient coordinately specifies sensory and support cell fates. (A-C) Representative images of the effect that fgfra knockdown has on support cell specification. (D) Schematic depicting the treatment regimen for treating embryos with the FGFR inhibitor SU5402. (E-K’) Double fluorescent in situ hybridizations visualizing sensory cells (prd146) and support cells (poxA) within the same larvae in control larvae (E, E’) or larvae treated with 10uM (G, G’), 15uM (I, I’), or 20uM (K, K’) SU5402 for 48 hours. (L) Schematic representation of how a gradient of FGF activity could coordinately specify both sensory and support cell fates. (M-O’) Double fluorescent in situ hybridizations co-staining prd146 and poxA in wild type (*fgfra* +/+), heterozygous (*fgfra^+^*^5^/+), and homozygous (*fgfra^+^*^5^*^/+^*^5^) mutant siblings. (P) Quantification of the circumference of the support cell ring in F2 *fgfra^+^*^5^ siblings. Initial differences among groups was determined by the non-parametric Kruskal-Wallis test (*H*(2) = 68.9, *p <* 0.0001, 17^2^_H_ = 0.5) and significant differences between groups was determined using Dunn’s post hoc test; ** *p.adjust* < 0.01, **** *p.adjust* < 0.0001 ****. Scale bar, 50um.

To further explore the role of FGF signaling during *Nematostella* apical organ development, we employed CRISPR mutagenesis to create an *fgfra* mutant allele carrying a 5bp insertion (*fgfra^+^*^5^). This insertion results in a frame shift with a premature stop codon truncating the protein prior to the first Ig-domain in the extracellular ligand binding domain. As expected from our knockdown experiments, F1 sibling in-crosses resulted in normal apical organ development in F2 wild type progeny and a complete loss of *prd146* and *poxA* expression in their homozygous mutant siblings (**Figure 7M, M’, O, O’**). Remarkably, in some heterozygotes we noted that the circumference of the support cell ring looked smaller (**Figure 7N, N’**), similar to what we observed following intermediate levels of FGFR inhibition (**Figure 7G, G’)** Indeed, the average circumference of the support cell ring of heterozygous planulae was statistically smaller than their wild type siblings (**Figure 7P**). Collectively, these results support a model wherein distinct thresholds of FGF signaling activity induce unique cell identities within the aboral domain of *Nematostella* larvae.

## Discussion

In the present study, we used molecular and genetic approaches to uncover an unexpected degree of complexity in the mechanisms governing apical organ development in *Nematostella vectensis*. Among marine invertebrates, relatively little is known about the cell type composition of larval apical sensory organs despite their prevalence and potential ecological importance. Indeed, most studies have focused on the role of apical organ-associated neuronal populations in regulating larval behavior and life stage transitions^11, 17, 19, 32, 63^. More recent work has sought to molecularly characterize the cell type populations associated with apical sensory organs. These studies show deep conservation among the genes expressed within the apical organ and aboral larval domain across species^29–31, 33^, but whether this corresponds to cell type conservation has remained unclear. Recently, bulk RNA-seq on oral and aboral domains of bisected *Nematostella* larvae compared the top aborally-enriched genes to a previously published single cell atlas to indirectly identify unique cell populations^31^. This approach successfully identified apical organ sensory cells, two subtypes of gland/secretory cell, aborally enriched neuronal populations, and a population of larval-specific neuron that encircles the sensory cells. The identification of these cell types represents a significant step towards building molecular definitions of apical sensory organ associated cell types.

Here, with single cell resolution, we identified distinct cell types found within the aboral domain and apical sensory organ. Our analysis identified neurons, secretory cells, and sensory cells, which is consistent with the previous report (**Figure 2B, B’, F, F’**)^31^. Interestingly, in our analysis neurons and secretory cells largely clustered together (**Figure 2A, B, B’**), which may be indicative of their shared developmental lineage^45^. Importantly, our sub-clustering analysis identified several additional aboral cell types not identified previously. A large proportion of the cells in the aboral domain correspond to embryonic ectodermal cells but we further identified a pool of progenitor cells, cnidocytes, and a unique cell cluster that was enriched with cells which had previously been classified as larval specific neurons (**Figure 2A, C, C’, D, D’, E, E’, G, G’)**. This definition was based on the observation that these cells express *poxA* and *slc26a6*, which are marker genes for a larval-specific neuronal cell cluster in a planula single cell atlas^31, 46^. However, we confirmed that these marker genes are expressed by two anatomically distinct cell populations: cells that ring the apical sensory organ as well as by cells scattered throughout the larval body column (**Figure 2G, G’, Figure S6B, B’**)^31^. We therefore tested whether these were indeed a single population of cells or two distinct cell types that expressed the same marker gene. Combined bioinformatic and functional analysis confirmed that these cells comprise two distinct cell populations and further led us to identify the marker gene *amt1,* which was specifically expressed by support cells (**Figure S6, S7**). Intriguingly, in the whole planula single cell dataset (**Figure S1**), support cells clustered within the apical sensory organ cell cluster. It was only after we subclustered aboral and sensory organ cells that the support cells formed their own unique population. With the increasing number of larval single cell data sets from different species, it will be interesting to perform similar sub-clustering experiments to determine whether the presence of a non-sensory support cell population is a common feature of apical sensory organs.

The precise developmental lineage from which cells of the apical sensory organ are derived is one major remaining question. Whether a subset of cells identified within our progenitor cell cluster represent *bona fide* apical organ progenitors is not clear. In *Nematostella*, the apical sensory organ does not appear to be related to any known neural lineage. The apical sensory organ forms its own unique cell cluster in single cell analyses, distinct from any known neuronal or neural progenitor cell population (**Figure S2A**)^45, 46^. Further, these cells lack expression of known neural genes^31, 46^. Regardless, rigorous lineage tracing studies aimed at interrogating progenitor cell sub-populations (*soxC^+^, soxB2^+^, nanos1^+^* progenitors) during apical organ development are needed. Alternatively, the apical sensory organ may not be derived from a single committed progenitor cell type. Instead, a pool of undetermined aboral ectodermal epithelial cells may become progressively specified as apical organ cells, similar to the development of cranial sensory placodes in chordates^64, 65^, by inductive signals, likely FGF ligands (**Figure 7**)^66^.

Our single cell RNA seq analysis also allowed us to identify a paired-type (PRD) homeodomain containing transcription factor as the most highly expressed marker gene in the sensory cell cluster (**Figure 2A, F, F’, Figure S2B, Supplemental Table 3**). This gene has been identified several times by computational means and phylogenetic analyses firmly place this gene within the PRD-like family of transcription factors. However, due to relatively poor statistical support, this gene does not seem to belong to a specific gene family (**Figure S4, Supplemental Table 4**)^31, 48–51^. Therefore, we continue to refer to this gene as *prd146*^50^. While the precise identity of *prd146* is ambiguous, it does form a strong orthology group (bootstrap=98) with homologues from other anthozoans (*Exaiptasia diaphana, Acropora digitifera, Stylophora pistillata*) (**Figure S4**), which suggests that this gene may be anthozoan specific. Although the precise function of this gene in these other species is not clear.

The precise function of the apical sensory organ in marine invertebrate larvae is poorly understood. It is thought that the apical sensory organ could regulate metamorphosis in part because it is a larval specific structure and because ablation of apical organ cells blocks the induction of metamorphosis in the nudibranch *Phestilla sibogae*^13^. However, this does not seem to be the case for all species^67^, suggesting that despite being a conserved larval feature it could have evolved species-specific functions. Interestingly, *prd146* mutant (**Figure S5N-P**) and morphant^31^ planulae lack the apical sensory organ but successfully undergo the larval to polyp transition. This suggests that in *Nematostella*, Prd146 and the apical sensory organ are dispensable for the polyp transition. However, the apical sensory organ could control the precise timing of the larval to polyp transition. Indeed, activation of GABA_B_ receptors delays the transition of planulae into polyps^68^. Therefore, GABAergic signaling from the apical organ sensory cells (**Figure 1C, D**) could fine tune developmental timing as opposed to activating or inhibiting the larval-polyp transition in a binary fashion.

During development of the *Nematostella* apical sensory organ, Prd146 is required to promote sensory cell specification and repress support cell fates (**Figure 4D-F, Figure S5G-I, Figure 6 B-C’**). However, the precise mechanisms through which Prd146 regulates this cell fate decision are still unclear but likely involves the combined inputs of Hox1 and SoxB1. Our data revealed a complex transcription factor network that regulates both sensory and support cell specification (**Figure 6**). Both Prd146 and SoxB1 function to repress support cell specification (**Figure 6B-D’, K**). However, *prd146* is a SoxB1 target gene (**Figure 4K, P**) therefore it is expected that *soxB1* knockdown would phenocopy *prd146* knockdown. In addition, we showed that Hox1 is required for support cell specification (**Figure 6E-K**). Interestingly, Hox1 is also required for specification of apical organ sensory cells^36^. Precisely how Hox1 promotes both sensory and support cell specification remains an interesting question.

Fibroblast growth factor signaling is required for the specification of both sensory and support cells during apical organ development (**Figure 4H, M, Figure 7A, B**). How one signaling pathway might specify two distinct cell types was not clear. During vertebrate development, FGF signaling can function in a classic morphogen gradient to induced discrete cell identities^38–40^. Using both pharmacological and genetic methods, we found that an FGF activity gradient coordinately specifies sensory and support cell fates (**Figure 7E-P**). Mechanistically, high FGF signaling activity induces the expression of Prd146, which in turn functions to promote sensory cell specification while repressing support cell fates (**Figure 4F, Figure 6C, C’**). At lower levels of FGF activity, Prd146 expression is not induced therefore those cells are specified as support cells (**Figure7I, I**). In the absence of FGF signaling, neither sensory nor support cells are specified (**Figure 7K, K’**). Despite these new insights, how and when the FGF signaling gradient is established remains unclear. Heparan sulfate proteoglycans (HSPF) play a crucial role in regulating the diffusion and receptor presentation of FGF ligands^38^. During *Nematostella* development, members of the HSPG family of proteins are important for aboral patterning and could play a role in establishing the FGF signaling gradient^69^.

---

**Supplemental Figure 1: Single cell RNA-seq analysis of whole *Nematostella* larvae.** (A) UMAP representation of larval Nematostella cell types. (B) Heat map of all marker genes enriched in larval cell types.

**Supplemental Figure 2: Sub-clustering analysis reveals cell type diversity in the aboral domain of larval *Nematostella*.** (A) Approach used to sub-cluster cells from the aboral domain (*six3/6*^+^, *foxQ2a*^+^) or apical sensory organ (*fgfa1*^+^/apical sensory organ cluster) of the whole larvae data set to generate Aboral/Sensory Organ UMAP. (B) Heat map depicting the expression levels of top 10 most highly expressed marker genes that define specific aboral cell populations.

**Supplemental Figure 3: *prd146* is specifically expressed by apical organ sensory cells.** (A) Dot plot showing that most known apical sensory organ genes in *Nematostella* are enriched in both sensory cells and support cells. (B-F) Visualization of cells in the Aboral/Sensory Organ single cell data set that express select genes. (B, C) *six3/6* and *foxQ2a* are expressed in multiple cell types but are repressed or lowly expressed in sensory cells. (D, E) *hox1* and *soxB1* are broadly expressed by aboral cell types including both sensory and support cells. (F) *prd146* is specifically expressed by only apical organ sensory cells.

**Supplemental Figure 4: Maximum-likelihood phylogeny reveals that *Nematostella* Prd146 encodes a PRD-like homeobox transcription factor that may be anthozoan-specific.** Predicted homedomains were aligned using L-INS-I via MAFFT; alignment file is shown in Supplementary materials. Gene tree was constructed with LG + G substitution model using RAXML with 500 rapid bootstraps. All bootstrap values are shown. The black box indicates a well-supported (bootstrap = 98) orthologous group containing NvPRD146 and three additional anthozoan sequences. Representatives of the MSX family of ANTP homeoboxes are used as outgroup. Species are color-coded as shown in figure. Abbreviations as followed: Nv = Nematostella vectensis; Ed = Exaiptasia diaphana; Spis = Stylophora pistillata; Aur = Aurelia coerulea; Clytia = Clytia hemisphaerica; Hydra = Hydra vulgaris; Hech = Hydractinia echinata; Ce = Caenorhabditis elegans; Pdum = Platynereis dumerilii; Spur = Strongylocentrotus purpuratus; Lvar = Lytechinus variegatus; Pmini = Patiria miniata; Skowl = Saccoglossus kowalesvskii; Bf = Branchiostoma florida; Hs = Homo sapiens.

**Supplemental Figure 5: Prd146 is required for apical sensory organ development but dispensable for the larval-polyp transition.** (A-F’) Fluorescent images of a developmental time course from Figure 3 specifically looking at Prd146 (magenta) expression dynamics, nuclei are in grey. (G-I) Fluorescent images showing the apical sensory organ in control (G) and *prd146* knockdown planula larvae using two independent shRNAs (H, I). (J-L) Immunofluorescent images showing Prd146 protein expression in control (J) and after *prd146* knockdown with two independent shRNAs (K, L). (M) Bar graph quantifying the roughly Mendelian ratio of the *prd146^+^*^4^ allele in 92 genotyped F2 primary polyps. (N-P) Actin staining showing the morphology of wild type (*prd146* +/+), heterozygous (*prd146^+^*^4^/+), and homozygous mutant (*prd146^+^*^4^/prd146*^+^*^4^) primary polyps. Scale bar, 15um (F’), 50um (A-E, G-L), 100um (F, N-P).

**Supplemental Figure 6: *slc26a6* and *poxA* are largely co-expressed in the same cell populations.** (A) Expression of *slc26a6* in the Aboral/Sensory Organ single cell data set. (B, B’) Fluorescent in situ hybridization of *slc26a6* in planula larvae. (C) Many cells that express *slc26a6* co-express *poxA* and are found within the Support Cell and Secretory Cells & Neurons clusters (black boxes).

**Supplemental Figure 7: The cells encircling the apical sensory organ are not neurons.** (A) Expression of *poxA* within the Aboral/Sensory Organ single cell data set is restricted to support cells and a subset of cells within the Secretory Cells & Neurons clusters (black boxes). (B) *poxA*^+^ cells within the Secretory Cells & Neurons cluster are likely neurons since they co-express the neuronal marker *elav* (black box). (C-D’) Inhibition of Notch signaling by DAPT treatment results in a loss of *poxA*^+^ cells encircling the apical sensory organ (compare C to D, C’ to D’; white boxes) but an expansion of the more lateral population of *poxA*^+^ cells (compare C to D; white arrow heads). (E) *amt1* expression is specifically enriched in cells of the Support Cell cluster (black box). (F) Only cells in the Support Cell cluster and not the Secretory Cells & Neurons cluster co-express *amt1* and *poxA*. (G-H’) Inhibition of Notch signaling by DAPT treatment results in a loss cells expressing amt1 that encircle the apical sensory organ (compare G, G’ to H, H’; white boxes).

## Acknowledgements

The authors want to thank members of the Gibson and Piotrowski labs for helpful comments as this project developed and Tak Akiyama for helpful comments on the manuscript. We also want to thank the Stowers Aquatics team for their dedicated care of our *Nematostella* colony. We also want to thank Sofia Robb and Whitney Leach for bioinformatics and computational support and well as Kym Delvanthal and Brandon Miller for guidance and team management for the CRISPR experiments and Jailyn Marshall for preparing Cas9 reagents. This work was funded by the Stowers Institute for Medical Research and the authors declare no competing interests.

## Methods

### Animal Husbandry

Adult Nematostella were housed in glass Pyrex dishes, kept largely in the dark, and cultured at 16°C. Animals were fed 2-5 times a week with freshly hatched brine shrimp and the morning before being spawned were hand fed mussel. Spawning was induced as previously described^70^. Briefly, bowls containing only females or a mix of males and females were placed in front of a light box at room temperature the evening before spawn induction and were exposed to light for 12 hours. The following morning, unfertilized egg gellies were collected from the female only dishes and were degellied with 4% cysteine solution in 12ppt artificial sea water for 10 minutes. After being washed with plain 12ppt ASW the eggs were kept at 16°C until injected with shRNAs (described below) and then fertilized with sea water from the mixed sex bowls. Fertilized eggs were kept at 24°C and fixed at 24hpf gastrula, 48hpf early planula, 72hpf planula, 96hpf late planula,120hpf tentacle bud, and 8dpf for primary polyp.

### Phylogenetic Analysis

Selected PRD sequences were obtained from the following 16 species: anthozoans *Nematostella vectensis*, *Exaptasia diaphana*, *Stylophora pistillata*, and *Acropora digitera*, medusozoans *Hydra vulgaris*, *Clytia hemispherica*, *Aurelia coerulea* (as *Aurelia* sp. 1), and *Hydractinia echinata*, and bilateria representatives both with apical organs, including *Strongylocentrotus purpuratus, Lytechinus variegatus, Patiria miniata, Platynereis dumerilii, Saccoglossus kowalevski*, and non-apical organs, including *Branchiostoma floridae, Caenorhabditis elegans*, and *Homo sapiens*. Candidate sequences were collected from either previously published studies and databases or NCBI searches using “homeobox” and the respective taxa. Sources for all sequences used in both datasets, including which sequences overall between the two, are provided in the supplementary materials. All sequences in both datasets were queried against the Pfam database (v35)^71^ using hmmsearch via HMMER (v3.3.2; http://hmmer.org/) with an E-value <= 0.001. The predicted homeodomains (PF00046.32) for all sequences were extracted and adjusted to include 60 amino acids as in previous studies^50^. The resulting homeodomain sequences were aligned using L-INS-I algorithm (--maxiterate 1000 —localpair) using MAFFT (v7.487)^72^. LG+G was determined to be the best substitution model using Prottest (v3.4.2)^73^. Maximum likelihood construction was conducted using RAXML (v8.2.12)^74^. Two independent runs were conducted with 25 random starting trees, and results were compared to determine the best tree. In addition, 500 rapid bootstrap replicates were performed for statistical support. Sources for all sequences are provided in the supplementary materials. As with Gilbert et al. (2022), sequences in the MSX family of ANTP class homeobox were used as an outgroup. All final trees were modified in FigTree (v1.4.4; http://tree.bio.ed.ac.uk/software/figtree/) and final figures produced using Inkscape (v1.2.1; https://inkscape.org/release/inkscape-1.2/).

### shRNA synthesis and embryo injection

Synthesis of shRNAs was performed as previously described^75, 76^. Briefly, a Universal Primer including part of the T7 promoter was mixed with a gene specific primer and annealed by heating to 70°C for 3 minutes before being allowed to cool to room temp for 5-10 minutes. The annealed primers were used as the template for a Klenow reaction, which ran for 30 minutes at 37°C, before the Klenow fragment was heat inactivated at 60°C for 20 minutes. This produced was used as the template for an T7 in vitro transcription kit which ran for 5-10 hours at 37°C. After DNase treating the IVT to degrade the DNA template the resulting shRNA was isolated using Direct-zol™ RNA Miniprep Plus Kit as per the manufacturers direction. The shRNA was eluted in 35uL of water and the concentration was determined using an Nanodrop.

Injection mixtures were generated by adding water, shRNA (final concentration 1000ng/uL), and FITC injection dye (2ug/uL). Control eggs were injected with a GFP shRNA which we have previously shown is an appropriate control shRNA^77^. A list of gene specific shRNAs is provided in **Supplemental Table 5**. Following injection, eggs were fertilized and kept at 24°C until the indicated developmental stage.

### Transmission electron microscopy (TEM)

For TEM analysis, embryos were prefixed with 2.5% Glutaraldehyde and 2% Paraformaldehyde in 50 mM Sodium cacodylate containing 1% Sucrose and 1mM Calcium chloride (pH 7.4). The samples were post fixed in 50mM Sodium cacodylate buffered 1%Osmium tetroxide, then in 0.5% Uranyl acetate. After dehydration with a graded Ethanol series, samples were infiltrated and embedded in Epon resin (EMS, Fort Washington, PA). Ultrathin (60-80 nm) sections were cut with a diamond knife and collected on single-slot copper grids. Grids were post-stained with 4% Uranyl acetate in 70% Methanol and Sato’s Triple Lead. Images were acquired on a FEI transmission electron microscope (Tecnai Bio-Twin12, FEI) at 80kV.

### Antibody Stains

Planula were fixed with 4% paraformaldehyde+0.05% glutaraldehyde for 2-4 minutes and then 4% paraformaldehyde for 1 hour at room temperature on a shaker. Following fixation, samples were washed 5 times with phosphate buffered saline (PBS) + 0.2% Triton-X and each wash was for 5 minutes. Then samples were incubated in blocking buffer (PBTx+5% normal goat serum+1% bovine serum albumin) for 1 hour. Primary antibodies were diluted into blocking buffer, rabbit anti-GABA (1:500) and mouse anti-acetylated tubulin (1:1000), and samples were incubated at 4°C overnight on a shaker. The following day, samples were subjected to 5x 20 minute washes with PBTx. After the washes, samples were incubated with secondary antibodies (goat anti-rabbit 568, goat anti-mouse 488; 1:500 in blocking buffer) at 4°C overnight on a shaker. The following day samples were subjected to 5x 20 minute washes before being incubated with the far red DNA dye SiR-DNA (1:1000 in PBTx) at 4°C overnight on a shaker.

Finally, samples were rinsed once in PBTx and then incubated in Scale A2 before being mounted and imaged. For the custom antibody raised against *Nematostella* Prd146, embryos, larvae, and polyps were fixed for 1 hour at room temperature on a shaker with a formic acid fixative solution (4% PFA, 4.8% Formic Acid, 25mM EGTA pH8.0, 100mM HEPES-KOH pH 7.5)^78^. The rest of the antibody staining procedure was identical to the one described above.

For EdU stains, developmental stages were incubated with 50uM EdU for 30 minutes before being rinsed 3 times with fresh 1/3 ASW. Samples were then fixed with the formic acid solution for 1 hour at room temperature before the EdU detection protocol was performed as per manufacturers suggestions (Thermo Scientific). Following EdU detection, the samples were processed for detection of Prd146 protein, as described above.

All samples were imaged using a Leica SP8 scanning laser confocal microscope or an Andor Dragonfly spinning disk confocal microscope. Single channel and composite images were made using Fiji and changes to brightness and contrast were applied to the entire image.

### Prd146 purification and antibody production

Full length *Nematostella vectensis* Prd146 (269 aa) was cloned into pMAL-C6T by Gibson Assembly, using a codon-optimized Geneblock purchased from IDT. Sequenced plasmid was transformed in to BL21 (DE3) E. coli for protein expression. Cells were grown in 4 L of Terrific Broth to an OD600 of 0.5, chilled in an ice-water slurry, and induced with 0.1 mM IPTG at 16 C for 16 hours. The IPTG-induced culture was harvested, cells were lysed by lysozyme treatment and sonication in lysis buffer (20 mM Tris, pH 8.0; 300 mM NaCl, 0.5 mM DTT, 1X cOmplete EDTA-free Protease Inhibitor Cocktail), and then centrifuged in a Type 45 Ti rotor at 35k x rpm for 60 minutes. All purification steps were performed on an AKTA Go FPLC instrument. The clarified lysate was applied wot a 5 mL HisTrap HP column, washed extensively in lysis buffer, and eluted with lysis buffer containing 250 mM imidazole. Peak fractions, selected based on the FPLC chromatogram, were pooled and diluted 10X with lysis buffer and immediately applied to a 5 mL MBPTrap HP column, washed extensively with lysis buffer, and eluted with lysis buffer containing 10 mM maltose. Peak fractions, as determined by SDS-PAGE, were pooled and buffer exchanged into 20 mM Tris, pH 8.0; 100 mM NaCl, 1 mM DTT, 0.5 mM EDTA. AcTEV protease (Invitrogen) was added at a ratio of 0.2 units/ug, and the mixture was incubated for 2 hours at 30C. Following cleavage, the mixture was passed over a 5 mL HisTrap column to remove free MBP and AcTEV protease. Purified Prd146 was concentrated to 0.2 mg/mL and sent to Pacific Immunology for rabbit polyclonal antibody production. To remove any potential MBP-reactive antibodies, serum from immunized animals was depleted using an affinity column containing purified MBP. Next, anti-Prd146 antibodies were purified from MBP-depleted serum using an MBP-Prd146 affinity column. The affinity purified antibodies were then tested for reactivity against Prd146 in prepared *Nematostella vectensis* specimens.

### Fluorescent In Situ Hybridizations

Samples at various developmental stages were fixed with 4% paraformaldehyde+0.2% glutaraldehyde for 2-4 minutes and then 4% paraformaldehyde for 1 hour at room temperature on a shaker. The samples were then washed 5 times for 5 minutes each wash in PBTx+DEPC before being gradually stepped into 100% methanol (25%, 50%, 75%, 100% PBTx+DEPC/methanol mixes). Samples were rinsed with 100% methanol before being stored at -20°C. Once ready, samples were bleached (methanol+3% hydrogen peroxide) for 1 hour under a light source and then gradually rehydrated (100%, 75%, 50%, 25% methanol/PBTx+DEPC). Samples were then washed 4 times for 5 minutes each wash with PBTx+DEPC and then digested with proteinase k (20ug/mL) for 10 minutes. Following protK digest, the samples were immediately post-fixed in 4% paraformaldehyde for 30 minutes before being washed 5 times for 5 minutes each wash with PBTx+DEPC. Samples were then incubated in a 1:1 mixture of PBTx:pre-hyridization buffer (lacking torula RNA and dextran) for 10 minutes at room temperature and then in 100% pre-hybridization buffer for an additional 10 minutes. Finally, the samples were incubated in hybridization buffer at 60°C overnight. The following day, the respective gene specific riboprobes (**Supplemental Table 5**) were diluted into hybridization buffer (1ng/uL or 0.5ng/uL) and then denatured by incubating at 80°C for 10 minutes. Probes were snap cooled by incubating on ice for 2 minutes before being brought up to the hybridization temperature. Samples were added to the respective riboprobes and incubated 60°C for 48 hours. After hybridization, samples were washed in pre-hybridization buffer and then gradually stepped into 2x SSC buffer. Then samples were washed 3 times in 0.2x SSC and each wash was for 20 minutes before being washed in a 1:1 mixture of 0.2x SSC/TNT buffer and then 100% TNT. Finally, samples were brought to room temperature and washed twice in TNT before being incubated in blocking buffer (TNT+5% sheep serum+1% Roche blocking reagent) for 1 hour. Anti-DIG Fab fragments or Anti-FITC antibody conjugated to POD were diluted 1:1000 in blocking buffer and samples were incubated over night at 4°C overnight on a shaker. The following day, samples were washed at least 5 times with TNT before the fluorescent signal was developed. To develop the fluorescent signal, samples were incubated for 30 minutes in Cy3-TSA diluted 1:50 in Diluent buffer (PerkinElmer). Finally, samples were washed at least 4 times for 15 minutes each was in TNT before fluorescent signal was checked. Samples were counter stained with the far red DNA dye, SiR-DNA, overnight at 4°C overnight on a shaker. The next day samples were rinsed with PBTx and then incubated in Scale A2 before being mounted and imaged.

For double fluorescent in situ hybridization, the samples were incubated with anti-FITC antibody conjugated to POD (1:1000) overnight, processed as described above, and the fluorescent signal was detected with Cy3. Then the peroxidase activity was quenched by incubating the samples in 200mM sodium azide in TNT buffer for 1 hour on a shaker at room temperature. The samples were washed 5x for 20 minutes each wash before being incubated in anti-DIG Fab fragments overnight at 4°C overnight on a shaker. The next day, samples were washed 5x with TNT buffer and the second florescent signal was detected by incubating the samples in Cy5-TSA diluted 1:50 in Diluent buffer (PerkinElmer). The samples were washed at least 4x for 15 minutes each wash and then stained with Hoechst at 4°C overnight on a shaker before being put into Scale A2.

### Single Cell RNA Sequencing and Analysis

Dissociated cells, having been sorted in PBS, were assessed for concentration and viability via a Nexcelom Cellometer Auto T4. Cells, having been deemed to be at least 65% viable, were loaded on a Chromium Single Cell Controller (10x Genomics, Pleasanton, CA), based on live cell concentration. Libraries were prepared using the Chromium Single Cell 3’ Library & Gel Bead Kit v2 (10x Genomics) according to manufacturer’s directions. Resulting short fragment libraries were checked for quality and quantity using an Agilent 2100 Bioanalyzer and Invitrogen Qubit Fluorometer. With cells captured estimated at ∼2,500 cells, the single cell library was sequenced to a depth necessary to achieve ∼135,000 mean reads per cell, or ∼330M reads total, on an Illumina HiSeq 2500 instrument, RTA v1.18.64, using Rapid SBS v2 chemistry with the following paired read lengths: 26 bp Read1, 8 bp I7 Index and 98 bp Read2. Raw reads were demultiplexed using 10X Genomics pipeline cellranger mkfastq. Genome and transcriptome indexes were built using Nematostella genome Nvec200. Fastq files were aligned by STAR aligner and cell expression table were generated using cellranger count function with default parameters. Cells with more than 500 UMI counts were loaded into analysis package Seurat for downstream analysis. Aboral and sensory subclusters were then extracted and re-clustered based on marker genes. Cell identities were assigned based on previously published marker genes^45^ (**Supplemental Table 1**) and our own experimental validation of cluster enriched gene expression.

### Prd146 and Fgfra CRISPR Mutagenesis and Genotyping

CRISPR-Cas9 technology was used to engineer knockout Nematostella vectensis strains. A guideRNA target site was selected using the CCTop target predictor tool^79^. The target site was selected by evaluating the predicted on-target efficiency score and the off-target potential^80^. The selected guideRNA was ordered as an Alt-R CRISPR-Cas9 sgRNA from Integrated DNA Technologies (IDT). The ribonucleoprotein (RNP) complex was formed with 500ng/ul sgRNA and 500ng/ul IDT Alt-R Cas9-GFP v3 protein and incubated at room temperature for 20 minutes. The RNP complex was delivered to *Nematostella* embryos by microinjection.

Tissue from resulting animals was lysed using QuickExtract DNA Extraction Solution (Epicentre) to release the genomic DNA. PCR was performed to amplify the specific genomic location, followed by a second round of amplification to incorporate sample-specific dual barcodes. All amplicons were pooled and size-selected using ProNex Size-Selective Purification System (Promega). Cleaned pools were quantified on a Qubit Fluorometer and then ran on an Agilent Bioanalyzer to check sizing and purity. Purified pools were run on an Illumina MiSeq 2x250 flow cell. The resulting sequence data was demultiplexed, and read pairs were joined. On-target indel frequency and expected mutations were analyzed using CRIS.py^81^.

### Drug Treatments

Eggs were fertilized as previously described and allowed to develop for 24-30 hours to ensure gastrulation was complete. Then embryos were incubated with 0.5% DMSO dissolved in 12ppt artificial sea water or 10uM, 15uM, or 20uM FGFR inhibitor (SU5402) or 10uM Notch inhibitor (DAPT) until 3dpf. The drug treatments were carried out in 6 well plates, drugs were changed every 24 hours, and were kept in the dark throughout the duration of the treatment.

Drug treated and control embryos were fixed at 3pdf for in situ hybridization as described above.

### Quantification of EdU^+^ and Prd146^+^ cells and Statistical Analyses

Images were imported into python. Every fifth slice of the z-stack, nuclei were segmented in the DAPI channel using Cellpose^82^ using cell diameter 9 and flow_threshold 0.4. Segmentation was checked for random samples at random depths into the tissue for accuracy. Three slices of IF channels around the segmented plane were max projected and the mean intensity was measured for each segmented cell through the z-stack. Cells were categorized as Prd146 or EdU positive using thresholds. The cell counts for each embryo were compiled and plotted in RStudio.

For statistical tests, data that did not meet the assumptions of homogeneity of variance, the non-parametric Kruskal-Wallis test was performed to determine if any differences existed among comparison groups. Dunn’s post hoc test was used to explore significant differences between the groups. The effect size for the Kruskal-Wallis analysis was determined using eta-squared based on the H statistic (ρι^2H^). A value of 0.01 indicates a small effect, a value of 0.06 indicates a medium effect, and a value of 0.14 or larger indicates a large effect^83^.

For data that met the assumptions for an ANOVA, an ANOVA was used to determine if any differences existed among comparison groups. Tukey’s HSD test was used to explore significant differences between the groups. The effect size for the ANOVA was determined using the generalized eta-squared (17^2^_G_). A value of 0.01 indicates a small effect, a value of 0.06 indicates a medium effect, and a value of 0.14 or larger indicates a large effect^83^.

An alpha level of 0.05 was used to determine statistical significance for all analyses. All analyses were performed, and graphs created, in R v 4.0.1 using the packages ggpubr and rstatix. Power calculations were done in G*Power v 3.1.9.2.

### Circumference x genotype results (Kruskal-Wallis) (n = 136 total, power ∼0.99)

Overall test: *H*(2) = 68.9, *p <* 0.0001, 17^2^_H_ = 0.5

Wild type vs Heterozygous: *z* = 3.4, *p.adjust* < 0.01 **

WT vs homozygous: *z* = 8.3, *p.adjust* < 0.0001 ****

Heterozygous vs Homozygous: *z* = 6.3, *p.adjust* < 0.0001 ****

### Percent of Prd146+ cells that are EdU+ (Kruskal-Wallis) (n = 55 total, power ∼0.99)

Overall test: *H*(2) = 40.6, *p <* 0.0001, 17^2^_H_ = 0.74

Gastrula vs. E.Planula: *z* = 3.3, *p.adjust* < 0.01 **

Gastrula vs. Planula: *z* = 4.7, *p.adjust* < 0.0001 ****

Gastrula vs. L.Planula-TB: *z* = 6.0, *p.adjust* < 0.0001 ****

E.Planula vs. Planula: *z* = 1.4, *p.adjust* = 1

E.Planula vs. L.Planula-TB: *z* = 2.8, *p.adjust* < 0.05 *

Planula vs. L.Planula-TB: *z* = 1.6, *p.adjust* = 0.6

## Notes

### Competing Interest Statement

The authors have declared no competing interest.

## References

1. A, K.-G., et al. Proliferation-independent regulation of organ size by Fgf/Notch signaling. eLife 6, (2017).

2. Chen, Y. et al. High-throughput sequencing of single neuron projections reveals spatial organization in the olfactory cortex. Cell 185, 4117–4134.e28 (2022).

3. Alsina, B. Mechanisms of cell specification and differentiation in vertebrate cranial sensory systems. Curr Opin Cell Biol 67, 79–85 (2020).

4. Alsina, B. & Whitfield, T. T. Sculpting the labyrinth: Morphogenesis of the developing inner ear. Semin Cell Dev Biol 65, 47–59 (2017).

5. Wang, S., Sengel, C., Emerson, M. M. & Cepko, C. L. A gene regulatory network controls the binary fate decision of rod and bipolar cells in the vertebrate retina. Dev Cell 30, 513–527 (2014).

6. Chen, Y.-C. & Desplan, C. Gene regulatory networks during the development of the Drosophila visual system. Curr Top Dev Biol 139, 89–125 (2020).

7. Young, C. M., Sewell, M. A. & Rice, M. E. Atlas of Marine Invertebrate Larvae. (Academic Press, 2002).

8. Nielsen, C. Trochophora larvae: cell-lineages, ciliary bands and body regions. 2. Other groups and general discussion. J Exp Zool B Mol Dev Evol 304, 401–447 (2005).

9. Nielsen, C. Trochophora larvae: cell-lineages, ciliary bands, and body regions. 1. Annelida and Mollusca. J Exp Zool B Mol Dev Evol 302, 35–68 (2004).

10. Conzelmann, M. et al. Conserved MIP receptor-ligand pair regulates Platynereis larval settlement. Proc Natl Acad Sci U S A 110, 8224–8229 (2013).

11. Conzelmann, M. et al. Neuropeptides regulate swimming depth of Platynereis larvae. Proc Natl Acad Sci U S A 108, E1174–1183 (2011).

12. Hadfield, M. & Paul, V. Natural Chemical Cues for Settlement and Metamorphosis of Marine-Invertebrate Larvae. in (eds. McClinTOCk, J. & Baker, B.) vol. 20015660 431–461 (CRC Press, 2001).

13. Hadfield, M. G., Meleshkevitch, E. A. & Boudko, D. Y. The apical sensory organ of a gastropod veliger is a receptor for settlement cues. Biol Bull 198, 67–76 (2000).

14. Katsukura, Y., David, C. N., Grimmelikhuijzen, C. J. P. & Sugiyama, T. Inhibition of metamorphosis by RFamide neuropeptides in planula larvae of Hydractinia echinata. Dev Genes Evol 213, 579–586 (2003).

15. Raff, R. A. Origins of the other metazoan body plans: the evolution of larval forms. Philos Trans R Soc Lond B Biol Sci 363, 1473–1479 (2008).

16. Page, L. R. Apical sensory organ in larvae of the patellogastropod Tectura scutum. Biol Bull 202, 6–22 (2002).

17. Kempf, S. C., Page, L. R. & Pires, A. Development of serotonin-like immunoreactivity in the embryos and larvae of nudibranch mollusks with emphasis on the structure and possible function of the apical sensory organ. J Comp Neurol 386, 507–528 (1997).

18. Magarlamov, T. Y., Dyachuk, V. & Chernyshev, A. V. Does the frontal sensory organ in adults of the hoplonemertean Quasitetrastemma stimpsoni originate from the larval apical organ? Front Zool 17, 2 (2020).

19. Miyamoto, N., Nakajima, Y., Wada, H. & Saito, Y. Development of the nervous system in the acorn worm Balanoglossus simodensis: insights into nervous system evolution. Evol Dev 12, 416–424 (2010).

20. Lacalli, T. C. The nervous system and ciliary band of Müller’s larva. Proc R Soc Lond B Biol Sci 217, 37–58 (1982).

21. Lacalli, T. C. & Berrill, N. J. Structure and development of the apical organ in trochophores of Spirobranchus polycerus, Phyllodoce maculata and Phyllodoce mucosa (Polychaeta). Proceedings of the Royal Society of London. Series B. Biological Sciences 212, 381–402 (1981).

22. LACALLI, T. C. Apical Organs, Epithelial Domains, and the Origin of the Chordate Central Nervous System1. American Zoologist 34, 533–541 (1994).

23. Voronezhskaya, E. E. & Khabarova, M. Y. Function of the apical sensory organ in the development of invertebrates. Dokl Biol Sci 390, 231–234 (2003).

24. Nakajima, Y., Burke, R. D. & Noda, Y. The Structure and Development of the Apical Ganglion in the Sea Urchin Pluteus Larvae of Strongylocentrotus droebachiensis and Mespilia globulus. *Development*, Growth & Differentiation 35, 531–538 (1993).

25. Nielsen, C. Larval nervous systems: true larval and precocious adult. J Exp Biol 218, 629–636 (2015).

26. Chia, F.-S. & Koss, R. Fine structural studies of the nervous system and the apical organ in the planula larva of the sea anemone Anthopleura elegantissima. J Morphol 160, 275–297 (1979).

27. Widersten, B. On the Morphology and Development in Some Cnidarian Larvae. (Almqvist & Wiksell, 1968).

28. Byrne, M., Nakajima, Y., Chee, F. C. & Burke, R. D. Apical organs in echinoderm larvae: insights into larval evolution in the Ambulacraria. Evol Dev 9, 432–445 (2007).

29. Marlow, H. et al. Larval body patterning and apical organs are conserved in animal evolution. BMC Biol 12, 7 (2014).

30. Sinigaglia, C., Busengdal, H., Lerner, A., Oliveri, P. & Rentzsch, F. Molecular characterization of the apical organ of the anthozoan Nematostella vectensis. Dev. Biol. 398, 120–133 (2015).

31. Gilbert, E. et al. Molecular and cellular architecture of the larval sensory organ in the cnidarian Nematostella vectensis. Development 149, dev200833 (2022).

32. Voronezhskaya, E. E., Khabarova, M. Y. & Nezlin, L. P. Apical sensory neurones mediate developmental retardation induced by conspecific environmental stimuli in freshwater pulmonate snails. Development 131, 3671–3680 (2004).

33. Feuda, R. & Peter, I. S. Homologous gene regulatory networks control development of apical organs and brains in Bilateria. Sci Adv 8, eabo2416 (2022).

34. Steinmetz, P. R. et al. Six3 demarcates the anterior-most developing brain region in bilaterian animals. EvoDevo 1, 14 (2010).

35. Rentzsch, F., Fritzenwanker, J. H., Scholz, C. B. & Technau, U. FGF signalling controls formation of the apical sensory organ in the cnidarian Nematostella vectensis. Development 135, 1761–1769 (2008).

36. Sinigaglia, C., Busengdal, H., Leclère, L., Technau, U. & Rentzsch, F. The bilaterian head patterning gene six3/6 controls aboral domain development in a cnidarian. PLoS Biol. 11, e1001488 (2013).

37. Amiel, A. R. et al. A bipolar role of the transcription factor ERG for cnidarian germ layer formation and apical domain patterning. Dev. Biol. 430, 346–361 (2017).

38. Balasubramanian, R. & Zhang, X. Mechanisms of FGF gradient formation during embryogenesis. Semin Cell Dev Biol 53, 94–100 (2016).

39. Yu, S. R. et al. Fgf8 morphogen gradient forms by a source-sink mechanism with freely diffusing molecules. Nature 461, 533–536 (2009).

40. Bökel, C. & Brand, M. Generation and interpretation of FGF morphogen gradients in vertebrates. Curr Opin Genet Dev 23, 415–422 (2013).

41. Wada, H. & Kawakami, K. Size control during organogenesis: Development of the lateral line organs in zebrafish. *Development*, Growth & Differentiation 57, 169–178 (2015).

42. Steiner, A. B., Kim, T., Cabot, V. & Hudspeth, A. J. Dynamic gene expression by putative hair-cell progenitors during regeneration in the zebrafish lateral line. Proc Natl Acad Sci U S A 111, E1393–1401 (2014).

43. Marlow, H. Q., Srivastava, M., Matus, D. Q., Rokhsar, D. & Martindale, M. Q. Anatomy and development of the nervous system of Nematostella vectensis, an anthozoan cnidarian. Dev Neurobiol 69, 235–254 (2009).

44. Kelava, I., Rentzsch, F. & Technau, U. Evolution of eumetazoan nervous systems: insights from cnidarians. *Philos. Trans. R. Soc. Lond., B*, Biol. Sci. 370, (2015).

45. Steger, J. et al. Single-cell transcriptomics identifies conserved regulators of neuroglandular lineages. Cell Rep 40, 111370 (2022).

46. Sebé-Pedrós, A. et al. Cnidarian Cell Type Diversity and Regulation Revealed by Whole-Organism Single-Cell RNA-Seq. Cell 173, 1520–1534.e20 (2018).

47. Zimmermann, B. et al. Sea anemone genomes reveal ancestral metazoan chromosomal macrosynteny. 2020.10.30.359448 Preprint at https://doi.org/10.1101/2020.10.30.359448 (2022).

48. Ryan, J. F. et al. The cnidarian-bilaterian ancestor possessed at least 56 homeoboxes: evidence from the starlet sea anemone, Nematostella vectensis. Genome Biol. 7, R64 (2006).

49. Chourrout, D. et al. Minimal ProtoHox cluster inferred from bilaterian and cnidarian Hox complements. Nature 442, 684–687 (2006).

50. Doonan, L. Transcriptome analysis in Hydractinia echinata reveals species-specific and phylum-wide gene loss. in (2018).

51. Mazza, M. E., Pang, K., Reitzel, A. M., Martindale, M. Q. & Finnerty, J. R. A conserved cluster of three PRD-class homeobox genes (homeobrain, rx and orthopedia) in the Cnidaria and Protostomia. EvoDevo 1, 3 (2010).

52. Pang, K., Matus, D. Q. & Martindale, M. Q. The ancestral role of COE genes may have been in chemoreception: evidence from the development of the sea anemone, Nematostella vectensis (Phylum Cnidaria; Class Anthozoa). Dev Genes Evol 214, 134–138 (2004).

53. Marlow, H., Matus, D. Q. & Martindale, M. Q. Ectopic activation of the canonical wnt signaling pathway affects ectodermal patterning along the primary axis during larval development in the anthozoan Nematostella vectensis. Dev. Biol. 380, 324–334 (2013).

54. Richards, G. S. & Rentzsch, F. Regulation of Nematostella neural progenitors by SoxB, Notch and bHLH genes. Development 142, 3332–3342 (2015).

55. Marlow, H., Roettinger, E., Boekhout, M. & Martindale, M. Q. Functional roles of Notch signaling in the cnidarian Nematostella vectensis. Dev. Biol. 362, 295–308 (2012).

56. Tournière, O., Gahan, J. M., Busengdal, H., Bartsch, N. & Rentzsch, F. Insm1-expressing neurons and secretory cells develop from a common pool of progenitors in the sea anemone Nematostella vectensis. Sci Adv 8, eabi7109 (2022).

57. Prelic, S. et al. Functional Interaction Between Drosophila Olfactory Sensory Neurons and Their Support Cells. Front Cell Neurosci 15, 789086 (2021).

58. Bucks, S. A. et al. Supporting cells remove and replace sensory receptor hair cells in a balance organ of adult mice. Elife 6, e18128 (2017).

59. Wan, G., Corfas, G. & Stone, J. S. Inner ear supporting cells: rethinking the silent majority. Semin Cell Dev Biol 24, 448–459 (2013).

60. Mohammadi, M. et al. Structures of the tyrosine kinase domain of fibroblast growth factor receptor in complex with inhibitors. Science 276, 955–960 (1997).

61. Marques, S. R., Lee, Y., Poss, K. D. & Yelon, D. Reiterative roles for FGF signaling in the establishment of size and proportion of the zebrafish heart. Dev Biol 321, 397–406 (2008).

62. Roehl, H. & Nüsslein-Volhard, C. Zebrafish pea3 and erm are general targets of FGF8 signaling. Curr Biol 11, 503–507 (2001).

63. Dickinson, A. J. G. & Croll, R. P. Development of the larval nervous system of the gastropod Ilyanassa obsoleta. J Comp Neurol 466, 197–218 (2003).

64. Koontz, A., Urrutia, H. A. & Bronner, M. E. Making a head: Neural crest and ectodermal placodes in cranial sensory development. Semin Cell Dev Biol 138, 15–27 (2023).

65. Saint-Jeannet, J.-P. & Moody, S. A. Establishing the pre-placodal region and breaking it into placodes with distinct identities. Dev Biol 389, 13–27 (2014).

66. Riley, B. B. Comparative assessment of Fgf’s diverse roles in inner ear development: A zebrafish perspective. Dev Dyn 250, 1524–1551 (2021).

67. Nedved, B. T., Freckelton, M. L. & Hadfield, M. G. Laser ablation of the apical sensory organ of Hydroides elegans (Polychaeta) does not inhibit detection of metamorphic cues. J Exp Biol 224, jeb242300 (2021).

68. Levy, S. et al. Ectopic activation of GABAB receptors inhibits neurogenesis and metamorphosis in the cnidarian Nematostella vectensis. Nat Ecol Evol 5, 111–121 (2021).

69. Bause, M., van der Horst, R. & Rentzsch, F. Glypican1/2/4/6 and sulfated glycosaminoglycans regulate the patterning of the primary body axis in the cnidarian Nematostella vectensis. Dev. Biol. 414, 108–120 (2016).

70. Stefanik, D. J., Friedman, L. E. & Finnerty, J. R. Collecting, rearing, spawning and inducing regeneration of the starlet sea anemone, Nematostella vectensis. Nat Protoc 8, 916–923 (2013).

71. Mistry, J. et al. Pfam: The protein families database in 2021. Nucleic Acids Res 49, D412–D419 (2020).

72. Katoh, K. & Standley, D. M. MAFFT Multiple Sequence Alignment Software Version 7: Improvements in Performance and Usability. Molecular Biology and Evolution 30, 772–780 (2013).

73. Darriba, D., Taboada, G. L., Doallo, R. & Posada, D. ProtTest 3: fast selection of best-fit models of protein evolution. Bioinformatics 27, 1164–1165 (2011).

74. Stamatakis, A. RAxML version 8: a tool for phylogenetic analysis and post-analysis of large phylogenies. Bioinformatics 30, 1312–1313 (2014).

75. Hill, E. M. et al. Manipulation of Gene Activity in the Regenerative Model Sea Anemone, Nematostella vectensis. Methods Mol Biol 2450, 437–465 (2022).

76. He, S. et al. An axial Hox code controls tissue segmentation and body patterning in Nematostella vectensis. Science 361, 1377–1380 (2018).

77. Karabulut, A., He, S., Chen, C.-Y., McKinney, S. A. & Gibson, M. C. Electroporation of short hairpin RNAs for rapid and efficient gene knockdown in the starlet sea anemone, Nematostella vectensis. Dev. Biol. 448, 7–15 (2019).

78. Guerrero-Hernández, C., Doddihal, V., Mann, F. G. & Alvarado, A. S. A powerful and versatile new fixation protocol for immunohistology and in situ hybridization that preserves delicate tissues in planaria. 2021.11.01.466817 Preprint at https://doi.org/10.1101/2021.11.01.466817 (2021).

79. Stemmer, M., Thumberger, T., Keyer, M. del S., Wittbrodt, J. & Mateo, J. L. CCTop: An Intuitive, Flexible and Reliable CRISPR/Cas9 Target Prediction Tool. PLOS ONE 10, e0124633 (2015).

80. Labuhn, M. et al. Refined sgRNA efficacy prediction improves large- and small-scale CRISPR–Cas9 applications. Nucleic Acids Research 46, 1375–1385 (2018).

81. Connelly, J. P. & Pruett-Miller, S. M. CRIS.py: A Versatile and High-throughput Analysis Program for CRISPR-based Genome Editing. Sci Rep 9, 4194 (2019).

82. Stringer, C., Wang, T., Michaelos, M. & Pachitariu, M. Cellpose: a generalist algorithm for cellular segmentation. Nat Methods 18, 100–106 (2021).

83. Vacha-Haase, T. & Thompson, B. How to Estimate and Interpret Various Effect Sizes. Journal of Counseling Psychology 51, 473–481 (2004).

