## Supplementary figures and images for "Graded FGF activity patterns distinct cell types within the apical sensory organ of the sea anemone *Nematostella vectensis*"

### Supplemental Figure 1

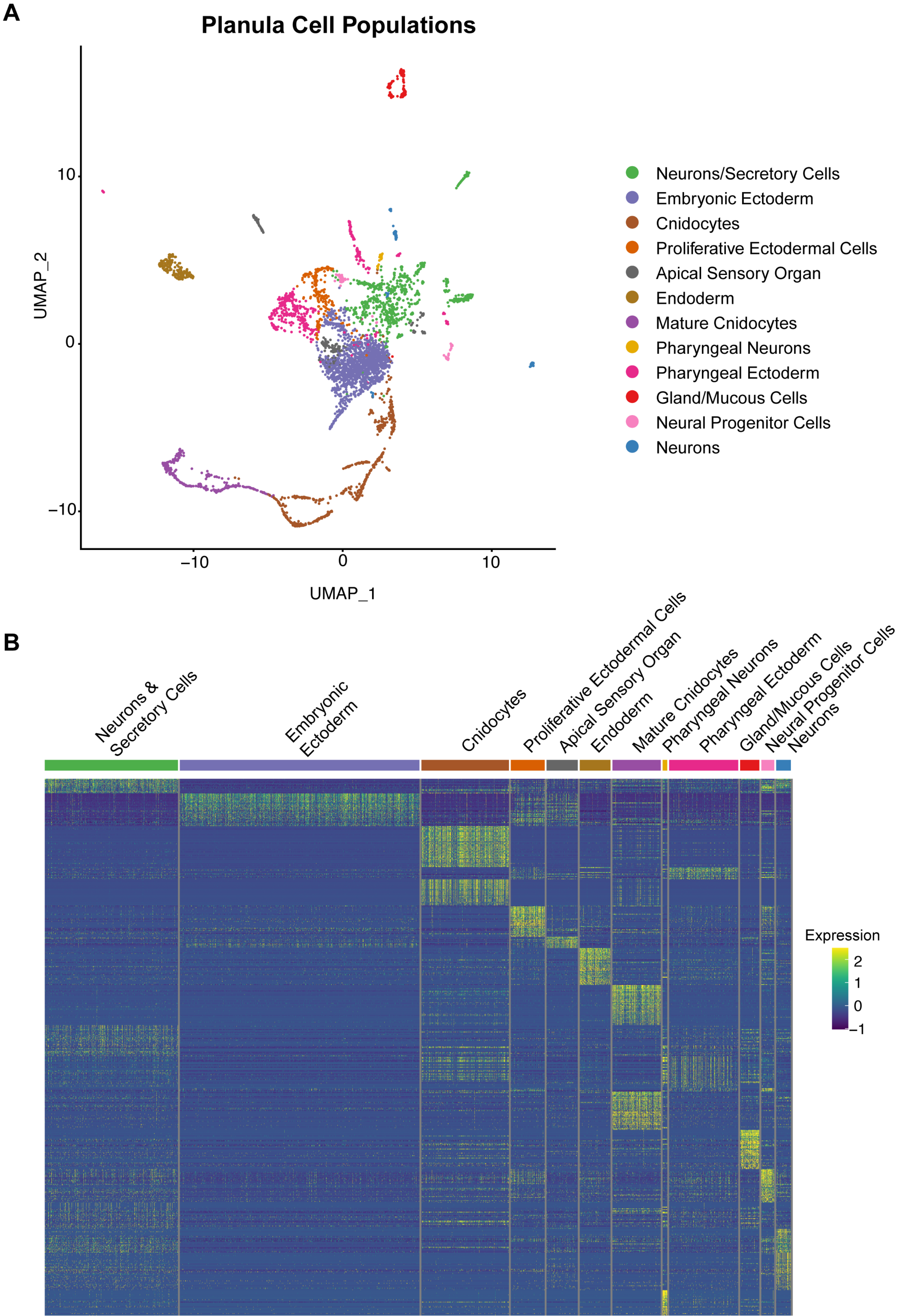

### Supplemental Figure 2

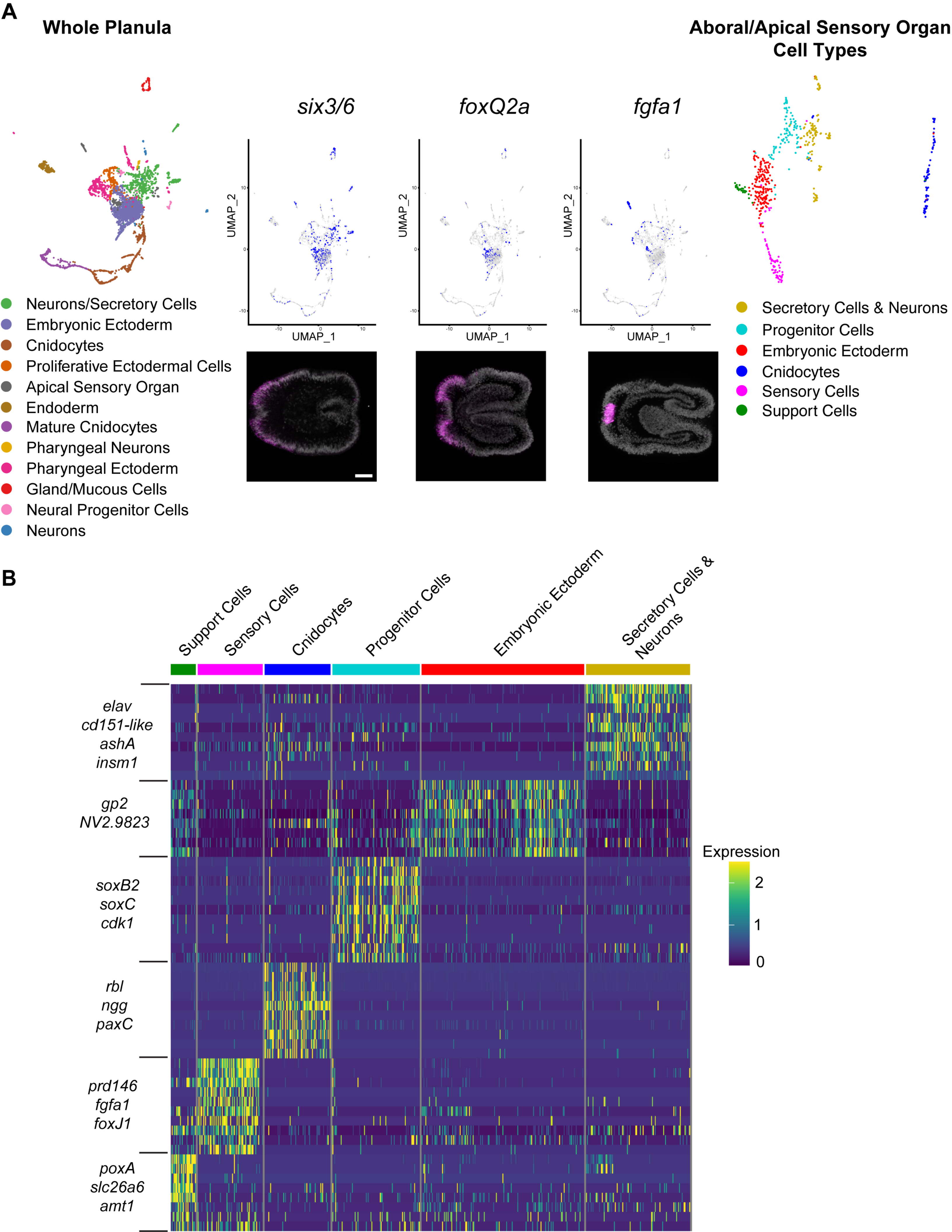

### Supplemental Figure 3

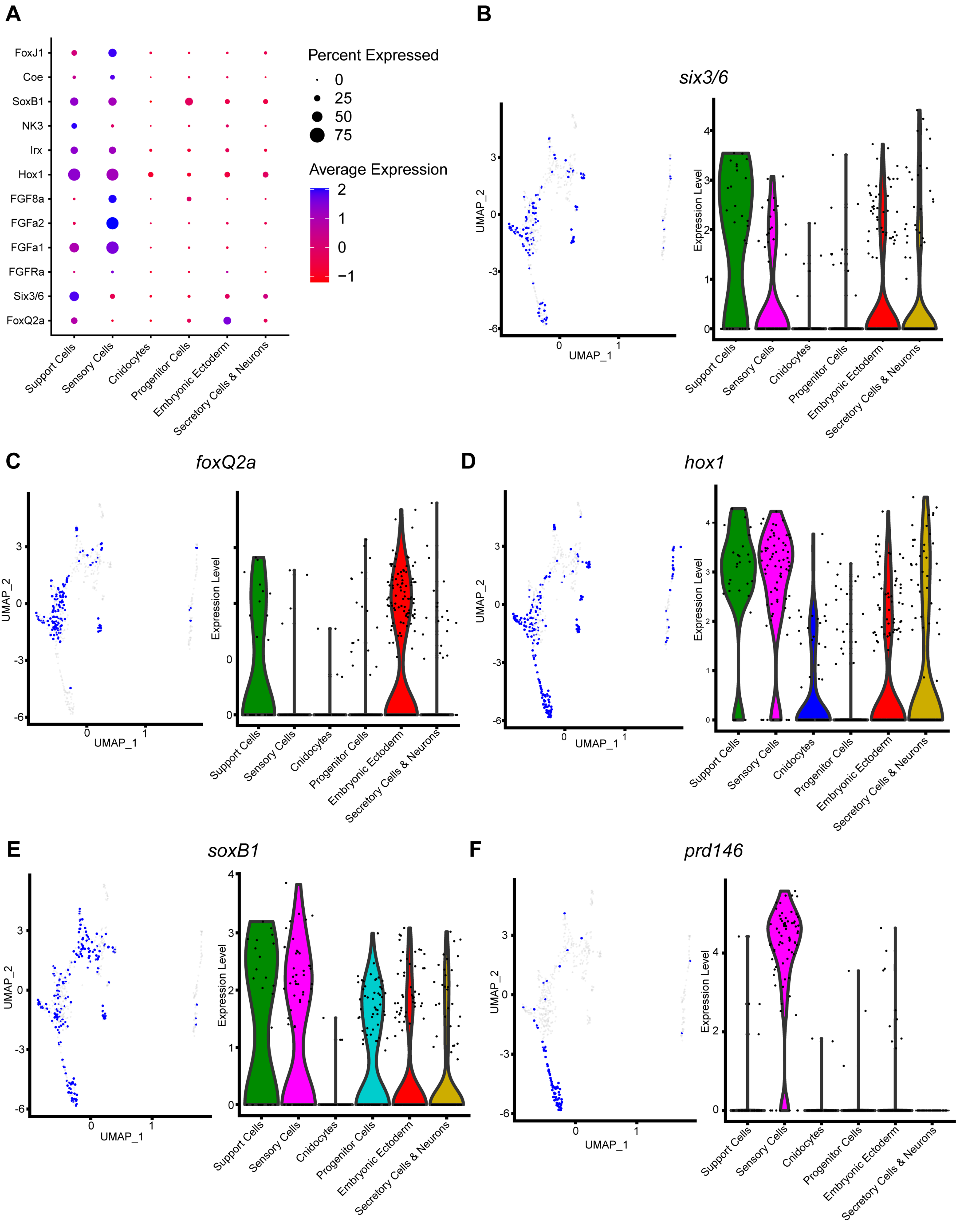

### Supplemental Figure 4

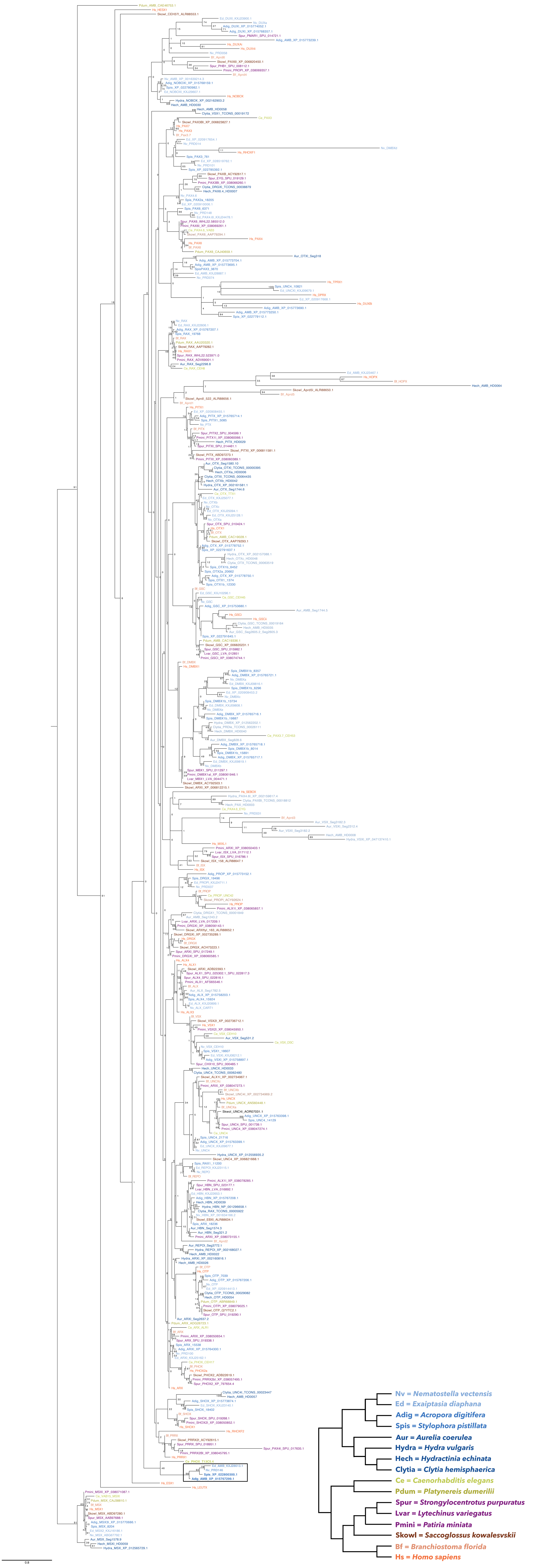

### Supplemental Figure 5

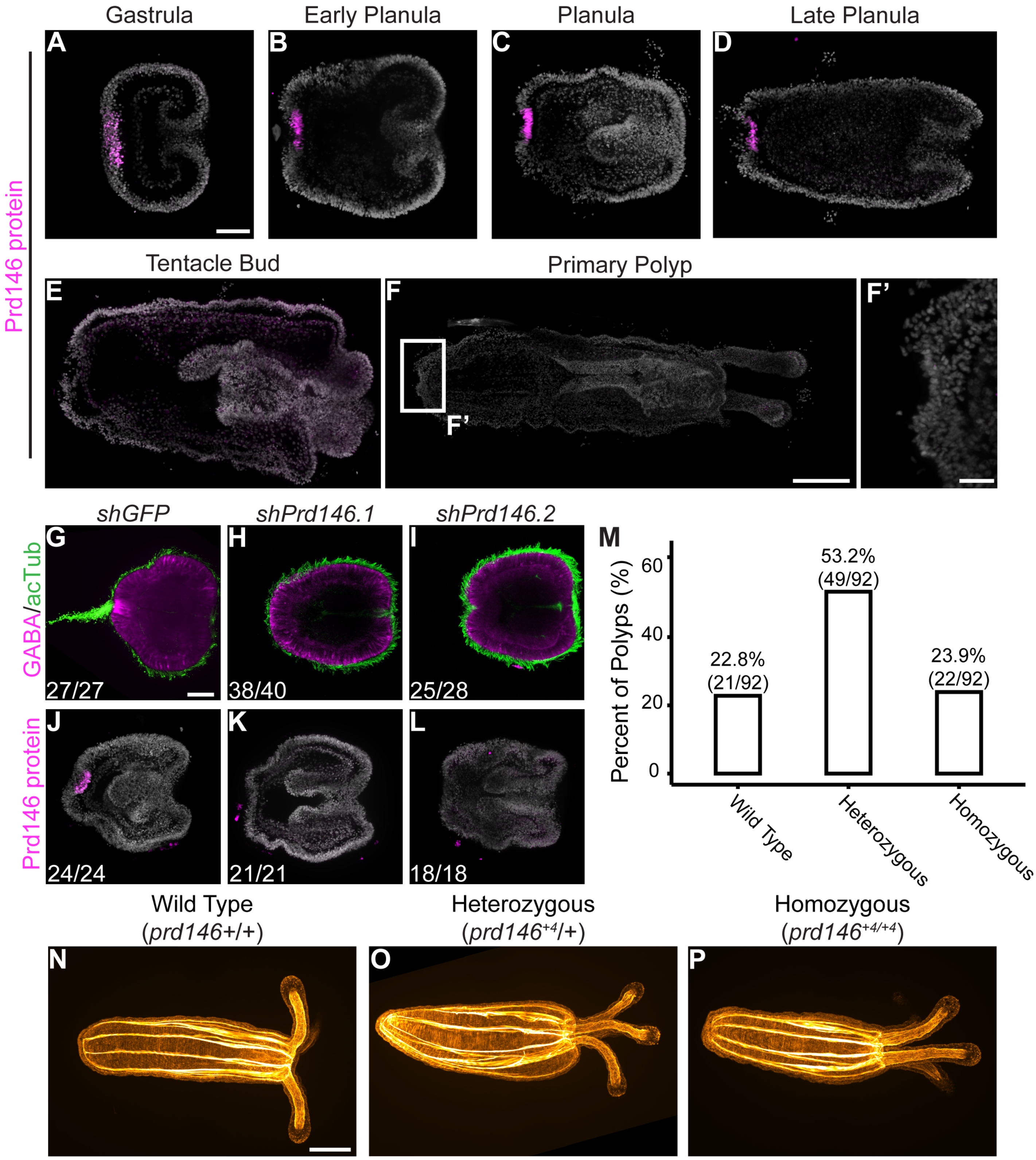

### Supplemental Figure 6

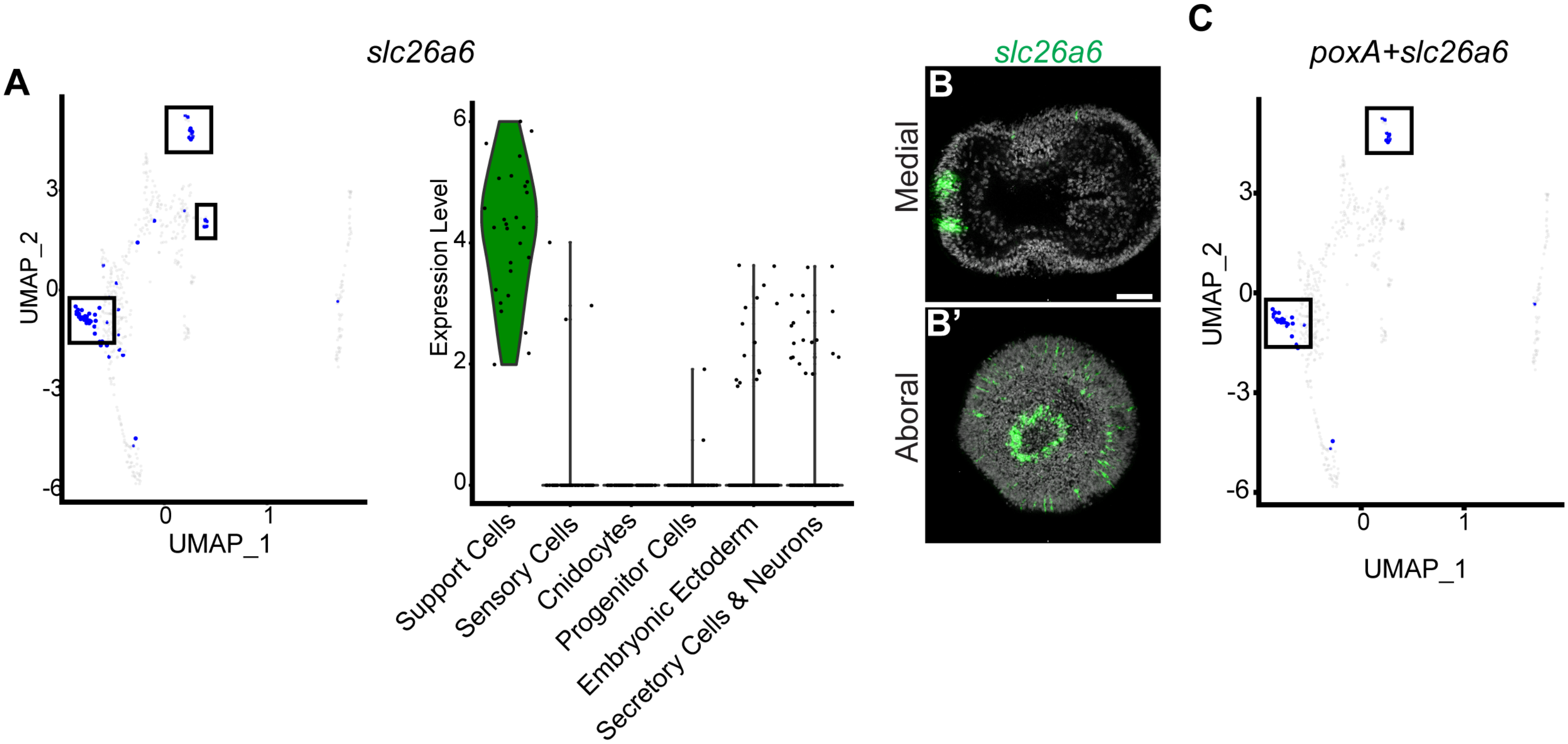

### Supplemental Figure 7

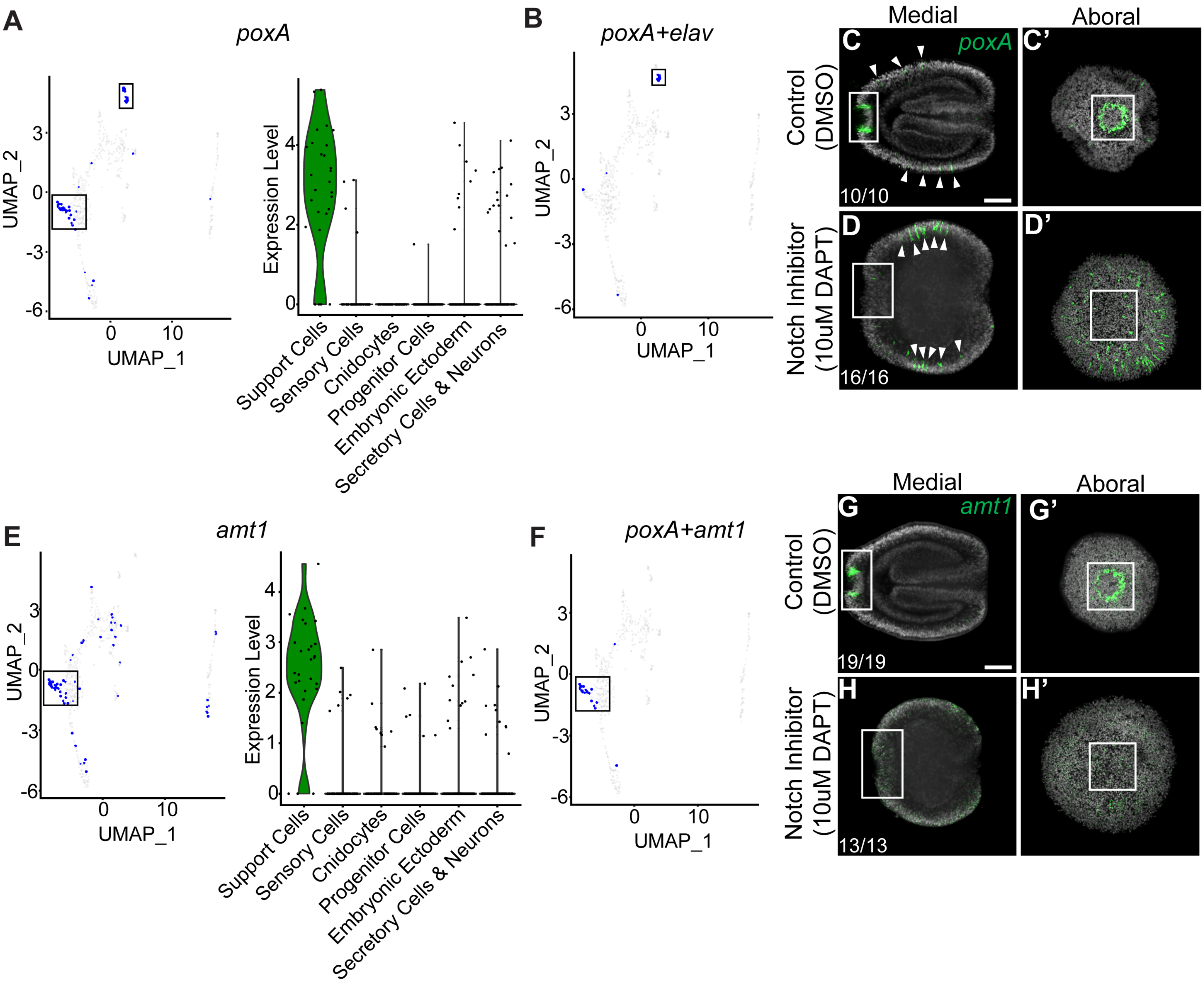
